# Test-Retest Reliability and Repeatability of Behavioral and Electrophysiological Markers in an Eriksen Flanker Task

**DOI:** 10.1101/2024.09.28.615624

**Authors:** Mario Bogdanov, Jason N. Scott, Shiba M. Esfand, Brian W. Boyle, Ty Lees, Mohan Li, Sarah E. Woronko, Samantha R. Linton, Courtney Miller, Shuang Li, Paula Bolton, Robert C. Meisner, Diego A. Pizzagalli

## Abstract

Cognitive control processes, specifically interference control and error monitoring, are often impaired across neuropsychiatric disorders and have been proposed as transdiagnostic markers of psychopathology and important treatment targets. Accurately probing them, however, requires understanding the psychometric properties of the measures used to assess cognitive control, including their intra- and interindividual stability over time. Using an Eriksen Flanker Task, we tested behavioral and electrophysiological readouts of cognitive control in 36 healthy individuals (26 female, 10 male, *M* age±*SD*=33.18±14.49, range=19–68) and evaluated their test-retest reliability across 48 hours by calculating Pearson correlations and Intraclass Correlation Coefficients (ICCs) to assess group-level stability. Moreover, we assessed repeatability through Coefficients of Variation (CVs) and Bland-Altman statistics, to investigate the degree of change in participants’ absolute scores. We found satisfactory-to-excellent test-retest reliability for most cognitive control measures, with condition-specific metrics generally being more reliable than difference scores. Regarding repeatability, we observed considerable intraindividual variability in absolute scores over time, which differed widely between participants. These results demonstrate that measurements of cognitive control may display substantial intraindividual variability across sessions despite demonstrating high test-retest reliability and vice versa. Our findings expand the current literature by providing novel information about the stability of behavioral and physiological markers of cognitive control over time. Moreover, they may have important implications for the application and evaluation of clinical interventions by highlighting the usefulness of considering repeatability measures in addition to the more commonly reported test-retest reliability metrics, when tracking changes over time in clinically relevant processes within single individuals.

## 1 Introduction

Interdisciplinary research has increasingly adopted multidimensional approaches that combine behavioral paradigms with neuroimaging or electrophysiological recordings in an effort to better understand mechanisms that characterize psychiatric disorders and identify processes that may serve as transdiagnostic markers of psychopathology (Montague et al., 2012; Niv, 2021; Pizzagalli et al., 2005; Wacker et al., 2009). For example, cognitive control, defined as the general ability to monitor and adjust thoughts and actions to enable goal-directed behavior, ranks among the processes most commonly impaired across psychiatric diagnoses (Goschke, 2014; Luciana & Collins, 2022; McTeague et al., 2016, 2017; Snyder et al., 2015), which led to its inclusion as a construct of interest in the Research Domain Criteria (RDoC) initiative (Insel, 2014). More specifically, a large number of studies reported reduced cognitive control (e.g., reduced interference control and performance monitoring) in individuals with depressive symptoms (Grahek et al., 2018; Romer & Pizzagalli, 2022), schizophrenia (Lesh et al., 2011), or substance use disorder (Ramey & Regier, 2019). Notably, these impairments have been linked to changes in neural processes that can be measured by electroencephalogram (EEG)-based techniques, such as event-related potentials (ERPs) and frequency band power (Clawson et al., 2013; de Aguiar Neto & Rosa, 2019; McLoughlin et al., 2022; Pathania et al., 2021; Ruchsow et al., 2004; Tenke et al., 2017; Vanderhasselt et al., 2012). In addition to identifying such maladaptive alterations at a given time, tracking changes in these cognitive and neural processes across time may provide a useful, more objective measure of the disease course or the efficacy of clinical interventions (McLoughlin et al., 2014; Olbrich & Arns, 2013; Riesel et al., 2021). However, the degree to which these measures may be useful toward this aim critically depends on their psychometric properties (Clayson, 2024; Cook & Beckman, 2006). To assess how an intervention affects cognitive processes, it is necessary to understand the expected variability of the behavioral or electrophysiological correlates of these processes over repeated measurements, for example by estimating their test-retest reliability. This is generally important in psychological research aimed at studying individual differences in cognition, but it is even more so in clinical trials aiming to evaluate novel treatments, given that samples in such studies are usually relatively modest in size and potentially heterogeneous with respect to the degree of cognitive impairment (Button et al., 2013; Schnack & Kahn, 2016).

Previous research has raised concerns about both the low frequency with which reliability estimates are reported and the actual degree of reliability displayed by commonly employed tasks (Enkavi et al., 2019; Hedge et al., 2018; Parsons et al., 2019; Vasey et al., 2003; Whitehead et al., 2019). For example, both cognitive (e.g., Stroop task) and affective (e.g., emotional dot-probe task) paradigms frequently show poor test-retest reliability, especially when compared to self-report measures that are far more often evaluated for their psychometric properties (Enkavi & Poldrack, 2021; Xu et al., 2024). This raises questions about whether these task-based behavioral measurements truly reflect stable, trait-like constructs, as would be required to draw meaningful conclusions about the clinical status of an individual or the possible effects of any specific treatment approach on cognition. Conversely, the necessity of testing and reporting measures of reliability is perhaps more common in electrophysiology (Clayson et al., 2019; Keil et al., 2014), where researchers often provide estimates of the psychometric qualities of their constructs of interest, such as ERP magnitudes and frequency band power. For example, numerous EEG studies have investigated the test-retest reliability of many common ERPs and associated frequency bands (Koo & Li, 2016; Larson et al., 2010; McEvoy et al., 2000; Morand-Beaulieu et al., 2021; Olvet & Hajcak, 2009; Tenke et al., 2017; Weinberg & Hajcak, 2011; Williams et al., 2005).

Across the literature, *test-retest reliability* is most commonly assessed by calculating correlation coefficients between multiple test sessions, such as a basic Pearson correlation or an Intraclass Correlation Coefficient (ICC), which account for systematic variability within participants and/or raters (Bartko, 1966; Koo & Li, 2016; Weir, 2005). Both approaches primarily quantify the degree of measurement error compared to the true interpersonal differences in the measure. Importantly, however, they reflect measurement stability primarily in terms of the strength of associations between testing sessions (i.e., it reflects stability of rank order or, whether people who score highest on a measure at one time also score highest on the measure at time two), not in terms of how similar a given participant’s measurement scores are in absolute values. To date, far less electrophysiological and behavioral research has systematically investigated this latter property, i.e., the *repeatability* of their measurements, which reflects the degree of within-person stability of a given measure. A common way of estimating repeatability is calculating Coefficients of Variation (CVs) – a unit-agnostic metric that assesses how similar repeated measurement scores from a single participant are across time, independent of the relative order of these measures between individuals (Aldridge et al., 2017; Berchtold, 2016; Duda et al., 2021; Hopkins, 2000; Shechtman, 2013; Vaz et al., 2013). However, CVs are meaningful only when applied to measurements on ratio scales with true zero-points, such as raw accuracy or RTs, but not on difference scores, such as the flanker interference effect (Shechtman, 2013). For measures on interval scales, Bland-Altman plots may serve as a fitting alternative (Bland & Altman, 1986, 1999). These plots depict average measurement scores across both assessments against the absolute difference between days as well as the 95% limits of agreement (i.e., mean difference ± 1.96SD) for said measure. This helps to visualize how well two measurements agree with each other (i.e., whether most individuals fall between the limits of agreement) and to highlight any systematic bias between them (i.e., whether the average difference between measurements significantly different from zero). Examining this intraindividual variability may provide useful information that can, in concert with test-retest reliability estimates, inform judgments about whether changes in any given measure (i.e., after patients have received a form of treatment) reflect meaningful improvements to the individual’s symptoms or simply expected fluctuations in these scores.

Thus, in this study, we aim to comprehensively examine the stability of behavioral and electrophysiologic indices of cognitive control over time as assessed by the Eriksen Flanker Task (Eriksen & Eriksen, 1974), a classic paradigm in which participants are required to respond to a centrally presented target stimulus while inhibiting responses to the surrounding congruent or incongruent distractors. We first investigated the test-retest reliability and repeatability of commonly reported behavioral measures of flanker task performance by assessing Pearson correlations and ICCs, and CVs and Bland-Altman statistics, respectively. More specifically, we considered accuracy and response times (RTs) in correct congruent and incongruent trials, as well as the flanker interference effect (i.e., the difference in both accuracy and RTs between congruent and incongruent trials; Eriksen & Eriksen, 1974; Eriksen, 1995), the Gratton effect (i.e., adjustments in accuracy and RTs in incongruent trials depending on the preceding trial type; Gratton et al., 1992), and the Laming and Rabbitt effects (i.e., post-error adjustments in accuracy and RTs, respectively; Laming, 1979; Rabbitt & Rodgers, 1977).

We then used the same approach to investigate the reliability and repeatability of several relevant ERPs and frequency bands previously linked to the cognitive control processes of inhibitory control and error monitoring. We examined the N200, a fronto-central ERP that is usually more negative for incongruent compared to congruent stimuli (Heil et al., 2000; Kanske & Kotz, 2010; Kopp et al., 1996), the error-related negativity (ERN), an early fronto-central ERP characterized by a pronounced negativity following erroneous responses compared to its counterpart following correct responses (CRN; Falkenstein et al., 1991; Gehring et al., 1993), and the error positivity (Pe), a late positive centro-parietal ERP following erroneous responses, as well as modulations in theta (4 – 7 Hz) and delta (0.5 – 4 Hz) power, given their well-established roles in cognitive control tasks (Cavanagh et al., 2012; Dell’Acqua et al., 2023; Karakaş et al., 2000; Muir et al., 2020; Sandre & Weinberg, 2019).

For all behavioral and electrophysiological measures, we expected to replicate typical findings in flanker tasks. Specifically, we predicted worse task performance (i.e., lower accuracy and slower RTs) and thus a larger flanker interference effect in incongruent vs. congruent trials, larger N200 amplitudes and theta and delta band power in response to incongruent vs. congruent targets, and larger ERN as well as Pe amplitudes and theta and delta power for erroneous vs. correct responses (Dell’Acqua et al., 2023; Falkenstein et al., 1991; Gehring et al., 1993; Heil et al., 2000; Kopp et al., 1996; Sandre & Weinberg, 2019). Moreover, we expected good-to-excellent test-retest reliability coefficients (Pearson r and ICCs), especially for condition-specific variables (e.g., accuracy, RTs, ERPs, and spectral power for congruent or incongruent trials, respectively), in line with previously reported reliability estimates for similar tasks and EEG signals (Morand-Beaulieu et al., 2021; Olvet & Hajcak, 2009; Tenke et al., 2017; Weinberg &

Hajcak, 2011; Williams et al., 2005). We also expected comparatively lower reliability for variables based on differences between conditions (e.g., the flanker interference effect, difference scores for ERPs and spectral power), given that simple difference scores usually yield smaller reliability estimates (Kessler, 1977; Miller & Kane, 2001; Paap & Sawi, 2016). Given that CVs have not yet been frequently used for cognitive or electrophysiological measures, we did not advance strong predictions about the range of these values. However, we assumed that raw behavioral measures (i.e., accuracy and RTs) might be more stable than measures that require more pre-processing steps (i.e., EEG power spectra). Finally, we predicted that differences in most measures would broadly center around zero, indicating good agreement across time, but that limits of agreement (i.e., the variability of these differences), based on Bland-Altman plots, would vary between measures.

## 2 Methods

### 2.1 Participants and experimental design

We recruited a total of 36 psychologically healthy individuals (26 female, 10 male, *M* age ± *SD* = 33.18 ± 14.49, range = 19 – 68) from the Greater Boston area. Participants self-identified as White (n = 22), Asian (n = 13) and Black or African American (n = 1); six participants identified as Hispanic or Latino. Participants did not report lifetime psychiatric disorders, as determined by the Mini International Neuropsychiatric Interview (MINI; Sheehan et al., 1998), which was administered by a masters or PhD-level clinical interviewer during a screening session. For some analyses, the sample was reduced due to insufficient performance or data for certain experimental conditions. We detail the reasons for exclusion and the final sample size used for a particular analysis below (see sections 2.3 and 2.4).

After passing the initial screening, participants visited the lab on two days, 48 hours apart, and completed two identical ∼2.5h long sessions in a within-subject design. On each day, participants first filled out a series of brief questionnaires before they were prepared for EEG and completed 8 minutes of resting baseline EEG (questionnaires and resting EEG are not relevant here and are not further discussed). This was followed by two experimental tasks—the Eriksen Flanker Task (see below) and a probabilistic reward task not relevant to the current study—in counter-balanced order. All participants provided written informed consent prior to study start and received monetary compensation (on average: $100 per session). Study procedures were conducted in accordance with the Declaration of Helsinki and were approved by the MGB Healthcare Institutional Review Board.

### 2.2 Eriksen Flanker Task

The Eriksen Flanker Task (Eriksen & Eriksen, 1974) is a widely used paradigm that probes cognitive control processes, specifically response inhibition, conflict monitoring and error processing. The task was displayed on a 22.5-inch VIEWPixx monitor (VPixx Technologies, Saint-Bruno, QC, Canada) using ePRIME software (ver. 2.0). Participants were seated approximately 70cm away from the monitor. In the task (see Figure 1), participants were presented with five horizontally displayed arrow stimuli (< or >) and were instructed to respond to the central arrow stimulus by pressing a button (left/right) on a response pad (RB-740, Cedrus Corporation, San Pedro, CA) that corresponded to the direction the arrow was pointing, while ignoring the direction of the flanking distractor stimuli. Distractor stimuli could either point in the same (congruent trials) or opposite (incongruent trials) direction as the central target arrow stimulus.

**Figure 1.**
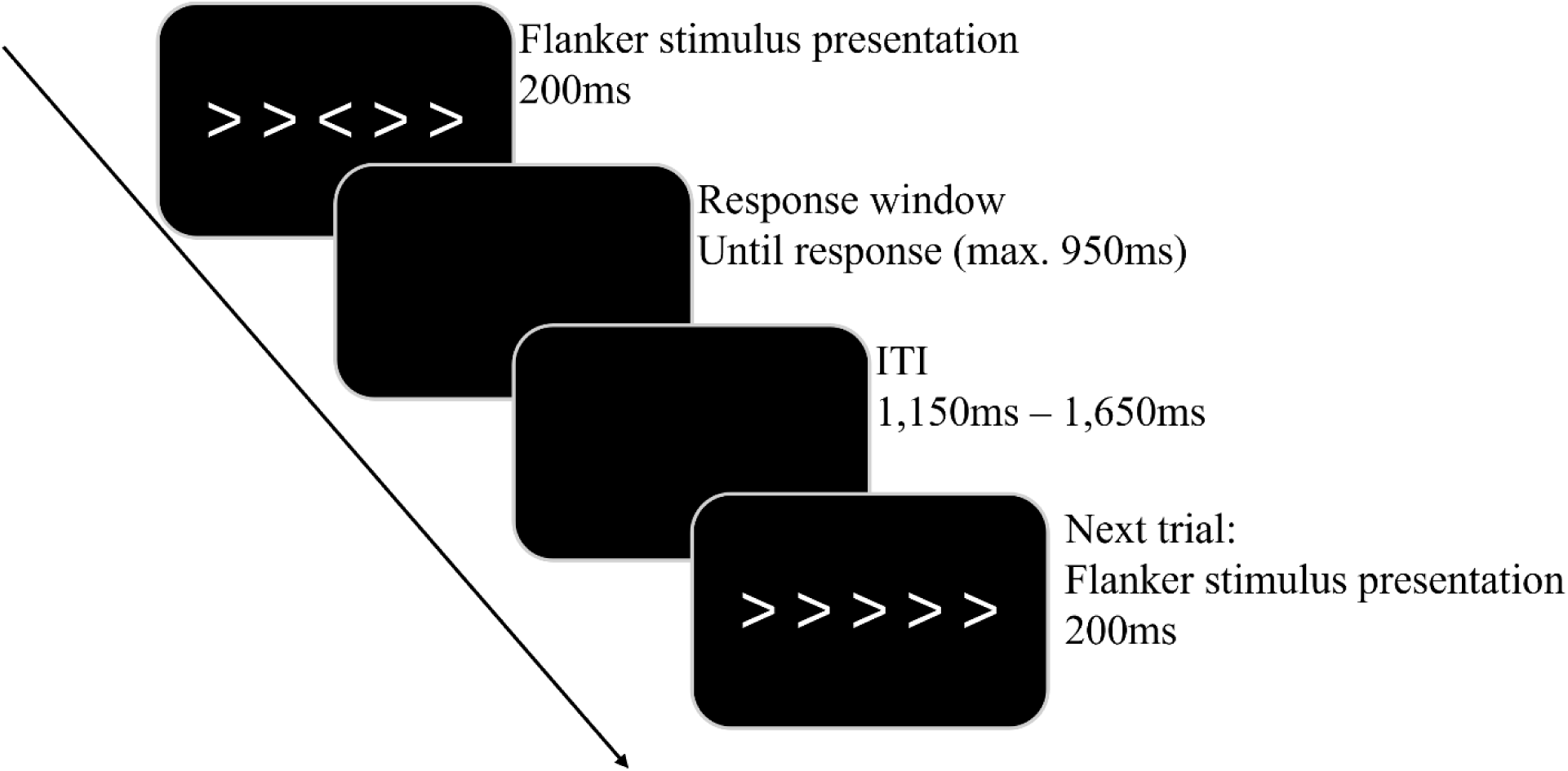
Eriksen Flanker Task. Participants had to indicate via button press whether the centrally presented arrow stimulus pointed to the left (<) or to the right (>) and to ignore the peripheral arrow stimuli (flankers). If the central arrow matched the direction of the flankers, the trial was “congruent”, and if they were pointing to different directions, the trial was “incongruent”.

In each trial, target and distractor stimuli (white on black background) appeared simultaneously on screen and were displayed for a total duration of 200ms. Then, a blank screen was presented for up to 950ms or until a response was recorded. This was followed by another blank screen with a random duration between 1,150 – 1,650ms, which served as the intertrial interval (ITI). The full task consisted of 240 trials (120 congruent/120 incongruent) equally presented across twelve blocks. In-between blocks, participants received general performance feedback by being presented with a screen showing the message “Please try to be more accurate” if average accuracy in the preceding block was below 75%, “Please try to respond faster” if average accuracy in the preceding block was above 90%, or “You’re doing a great job” if average accuracy was between these thresholds. The reason for the upper accuracy threshold of 90% was that we wanted to ensure participants produced sufficient incorrect responses to analyze behavioral and electrophysiological correlates of error processing (i.e., Laming/Rabbitt effect, ERN, Pe). On both testing days, participants completed a brief practice block prior to the main task in order to familiarize themselves with the paradigm. In total, it took participants ∼15min to complete the task.

### 2.3 Preparation of behavioral data

Behavioral output files provided by ePRIME were pre-processed using a custom Python (ver. 3.9) script. Preprocessing included identification of trials with missing responses (571 trials, 2.0% of data across all participants) and response times outside a pre-defined response window (200 – 1000ms; 288 trials, 1.0% of data across all participants). After a series of quality checks following pre-determined accuracy and response time criteria, we removed one participant from all behavioral analyses due to missing 32 (13.3%) and 42 (17.5%) of their trials on testing days 1 and 2, respectively (pre-determined cut-off: 24 trials, i.e., 10%). For analyses involving error responses (specifically the Laming/Rabbitt effects), we additionally excluded participants (*n* = 14) who committed too few (i.e., < 6) error responses on one or both of their testing days (Olvet & Hajcak, 2009), leading to a sample size of n = 22 in these analyses, to keep in line with our EEG-related exclusion criteria (detailed further below).

Following data cleaning, flanker interference effects were calculated for both accuracy (congruent – incongruent trials) and RTs (incongruent – congruent trials; correct responses only), separately for each testing day. For the Gratton effect, we identified incongruent trials that were preceded by either congruent or incongruent trials and calculated the difference in both accuracy (post incongruent – post congruent) and RTs (post congruent – post incongruent). Finally, for the Laming/Rabbitt effects, we identified incongruent trials that followed either a correct or an incorrect response to the previous trial and calculated the difference in both accuracy (post error – post correct) and RTs (post error – post correct).

### 2.4 EEG recording and data processing

#### 2.4.1 Data acquisition and general preprocessing

Participants completed the EEG data collection inside an acoustically and electrically shielded booth. Continuous EEG activity during the flanker task was recorded from a customized equidistant spherical 96-channel actiCAP system using an actiCHamp amplifier (Brain Products GmbH, Gilching, Germany). The ground electrode was embedded in the cap and was located anterior to the right of channel 10 which roughly corresponds to electrode AFz, and Channel 1 (Cz) served as the online reference. Data were digitized at a sample rate of 500Hz using BrainVision Recorder software (ver. 1.25.2001; Brain Products) with electrode impedance maintained below 25kΩ.

Data preprocessing was conducted offline using BrainVision Analyzer software (ver. 2.2; Brain Products). First, unsegmented data were visually inspected for muscle artifacts which were manually removed along with EEG data recorded during task breaks. Then, data were band-pass filtered using a zero-phase Butterworth filter with cutoffs set at 0.1 and 30 Hz, and a 12 dB/oct roll-off. Independent component analysis (ICA) was used to remove eye blinks, horizontal eye movements, and electrocardiogram, and corrupted channels were interpolated using spline interpolation. Lastly, data were re-referenced to the average activity of all electrodes. Pre-processed data were then segmented into 3,000ms epochs beginning 1,500ms before and 1,500ms after flanker stimulus onset (“target-locked” analyses) or response onset (“response-locked” analyses). Subsequently, automatic artifact rejection was performed so that intervals were rejected from individual channels in each trial if they contained a single voltage step exceeding 50μV, a voltage difference of more than 150μV in a 200ms interval, an amplitude of < −75μV or > 75μV within a trial, or a maximum voltage difference of less than 0.5μV in a 100ms interval. Further, in line with our behavioral criteria (see above), we rejected segments if responses fell outside our pre-determined response windows, i.e., if they were made too fast (< 200ms) or too slow (> 1,000ms).

#### 2.4.2 ERP Analyses

In total, we investigated three different ERP components. For all three components, choices for measurement windows and electrodes were based on previous research as well as a hypothesis- and data-independent collapsed localizer approach across all trial/response types and testing days (Luck & Gaspelin, 2017). Separate averages were created for task-specific conditions (i.e., congruent vs. incongruent trials for the N200, correct vs. incorrect responses for the ERN/CRN and the Pe) and testing days.

For the N200, the time from −200 to 0ms prior to stimulus onset served as the baseline, and the component was scored as the average activity at channel 2 (approximating electrode FCz) between 250 – 290ms following presentation onset of the flanker stimulus. Data from two participants were excluded from the target-locked analysis, one for failing to respond to a substantial number of trials on both days and a second participant due to technical errors with their EEG recording on day 2.

In turn, the ERN, CRN, and Pe were baseline-corrected using the time −800 to −700ms prior to responses. The ERN and CRN were then scored as the average activity at channel 9 (approximating Fz) between 10 - 50ms after response, while the Pe was scored as the average activity at Channel 1 (approximating Cz) between 160 - 200ms after response. For the error-related components, 12 participants were excluded for having too few error trials (< 6) after pre-processing. This cut-off was implemented to ensure sufficient trial counts for robust ERNs and is in line with prior recommendations (Olvet & Hajcak, 2009). For the remaining participants, we retained an average of 14.70 (SD = 6.88, range: 6 - 38) and 14.39 (SD = 7.60, range: 6 - 37) of artifact-free ERP segments for error responses on day 1 and 2, respectively, and an average of 99.13 (SD = 8.79, range: 69 - 110) and 99.87 (SD = 10.82, range: 67 - 113) of artifact-free ERP segments for correct responses on day 1 and 2, respectively.

#### 2.4.3 Time-Frequency Analyses

Time-frequency analyses for each time-window of interest were performed on the grand average for each task condition and testing day, with the time from −500 to −300ms prior to stimulus (N200 time-window) or response onset (ERN/CRN and Pe time-windows) used as the baseline, respectively. We performed a continuous wavelet transform using a Morlet complex with frequency bounds between 0.01 and 20Hz, 40 frequency steps, a Morlet parameter of 3.5 and Gabor normalization. We then extracted a single layer for each frequency band (central frequencies ± bandwidth: theta = 6.21 ± 3.55; delta = 2.85 ± 1.63) for each condition and testing day (Bogdanov et al., 2022). Absolute power in each frequency band was scored as the average activity at channels 2 (220 – 320ms), 9 (0 – 100ms) and 1 (130 – 230ms, which overlap with the time windows chosen for the respective ERP components (i.e., N200, ERN/CRN and Pe), but are extended on both directions to capture sufficient cycles of each frequency of interest for the wavelet decomposition of the signal. Data exclusions were identical to those reported for the corresponding ERP analyses.

### 2.5 Data analysis and reliability estimation

All inferential and reliability analyses were performed in R (ver. 4.2.2) using RStudio (ver. 2022.12.0). To verify successful implementation of response conflict, task-specific differences across testing days in behavioral (accuracy, RTs in correct trials) and electrophysiological measures (ERPs, spectral power), we performed 2 (trial/response type: “congruent” for vs. “incongruent” for the N200 or “correct” vs. “incorrect” for the ERN/CRN and Pe) × 2 (testing day: 1 vs. 2) repeated-measures Analyses of Variance (ANOVAs; Type III), with critical *p*-values set to *p* < .05. As an additional measure of data quality, we calculated split-half reliabilities (split by odd vs. even trials) for behavioral and EEG-derived indices within a single testing day using the Spearman-Brown correction (Hajcak et al., 2017; Pronk et al., 2022):

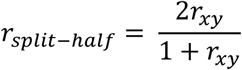

where r_xy_ is the Pearson correlation between both halves of each testing day.

For our main goal of assessing test-retest reliability and repeatability of these measures, we calculated several complementary indices. First, we calculated standard Pearson correlations (*r*), which estimate the strength of the relationship between the same measures across both days without accounting for any systematic variance caused by repeated measures. Then, we calculated two commonly used versions of the Intraclass Correlation Coefficient (ICC), which slightly differ with respect to their assumptions regarding sources of variance that contribute to the measurement error across sessions. In particular, we estimated ICC_3_, in which testing days are considered as fixed effects only and which focuses more on estimating overall consistency in the rank-order of individual measures across days, and ICC_2_, in which testing days are considered as random effects and estimates reliability with a larger focus on absolute agreement of individual values (instead of just relative order) across days, allowing for better generalization of reliability beyond the current study (Bartko, 1966). Although ICC_2_ and ICC_3_ are usually in close agreement, the former is conceptually closer to a measure of repeatability than the latter (Lin et al., 2020). Both ICCs were calculated using the ICC function from the R library psych (ver. 2.4.1). For *r* and both types of ICCs, we interpret coefficients as follows: < 0.6 poor; 0.6 – 0.8 acceptable; 0.8 – 0.9 good; > 0.9 excellent. Note that ICCs are theoretically constrained between 0 and 1 and that the negative estimates likely reflect low between-subject variance. Negative ICCs can be treated as factually zero (Bartko, 1976), however, we still included the estimated values in these cases for transparency. Finally, we estimated the Coefficient of Variation (CV) as a measure of repeatability. CVs were calculated using the following formula:

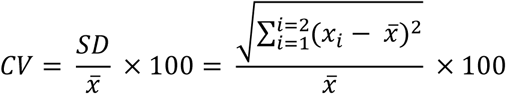

where 𝑥_𝑖_ is the value of the measure of interest on day *i* and SD refers to the standard deviation of that measure across both days (Duda et al., 2021; Shechtman, 2013). CVs were obtained for each participant separately. In short, CVs represent a percentage-wise measure of how much the exact values of a variable (e.g., RTs in error trials in day 1 and day 2) differ from the participant-specific average across days, with lower values suggesting higher repeatability. Importantly, the CV ignores whether the rank-order of participants stays consistent across time and purely assesses absolute score variation within participants. There are no established thresholds for which CVs can be considered adequate, given that these coefficients depend on the measure to which they are applied. In medical imaging, some have interpreted CVs of < 5% as very good, CVs of 5 – 10% as good, CVs of 10 – 20% as acceptable and CVs of > 20% as questionable (Aronhime et al., 2014); however, given the inherent variability in cognitive control processes (Braver, 2012), we expected CVs for behavioral and EEG-related measures to possibly be larger than these suggested ranges. We thus refrain from classifying these indices in terms of quality thresholds. As CVs can only be meaningfully interpreted for measures on ratio scales (Shechtman, 2013), we further created Bland-Altman plots and associated statistics as additional markers for repeatability using the BlandAltmanLeh package in R (Lehnert & Lehnert, 2015). An important aspect of repeatability concerns whether the average differences in a given measure between testing days center around zero, implying that there is no systematic bias between sessions. This is captured by our ANOVA analyses (see above), specifically in the main and interaction effects involving testing day for each measure. Further, we calculated the 95% limits of agreement that mark the mean between-day difference score ± 1.96SD for each measure and help to highlight the variance in the sample (i.e., how big is the range of values that lie within the limits) and the proportion of individuals that fall outside the limits. All figures were created with ggPlot2 (Wickham, 2011).

## 3 Results

### 3.1 Behavior

As expected, participants responded more accurately (*F*_(1, 34)_ = 68.92, *p* < .001, η^2^_G_ = 0.44) and faster (*F*_(1, 34)_ = 267.81, *p* < .001, η^2^_G_ = 0.30) on congruent trials compared to incongruent trials, demonstrating a robust flanker interference effect in both variables of interest (see Figure 2). While accuracy was comparable across both days (*F*_(1, 34)_ = 0.18, *p* = .675, η^2^_G_ < 0.01), participants’ response times were faster on the second day of testing (*F*_(1, 34)_ = 37.82, *p* < .001, η^2^_G_ = 0.05). We did not observe any significant trial type × day interaction effects (accuracy: *F*_(1, 34)_ = 0.01, *p* = .947, η^2^ < 0.01; RTs: *F*_(1, 34)_ = 3.76, *p* = .061, η^2^ < 0.01).

**Figure 2.**
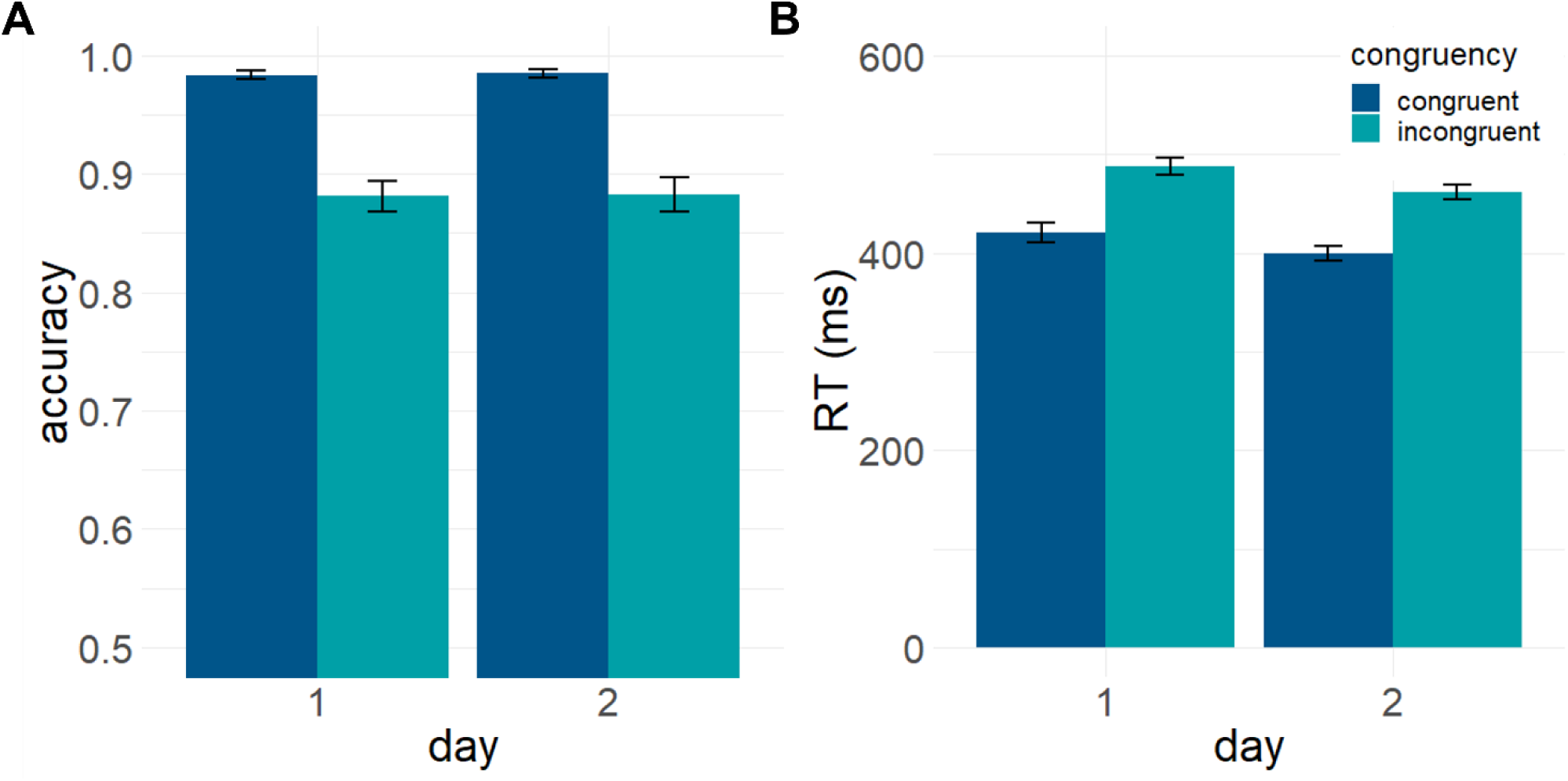
Behavioral task performance. Participants displayed the expected flanker interference effects. Specifically, accuracy (A) was significantly higher, and RTs (in correct trials; B) were significantly faster in congruent compared to incongruent trials on both days. While average accuracy was consistent across days, participants responded generally faster on day 2.

Further, we found that accuracy increased for incongruent trials that were preceded by incongruent compared to congruent trials (“Gratton effect”; *F*_(1, 34)_ = 47.72, *p* < .001, η^2^ = 0.06) on both testing days (main effect day: *F*_(1, 34)_ = 0.12, *p* = .723, η^2^ < 0.01; preceding trial type × day interaction: *F*_(1, 34)_ = 0.39, *p* = .535, η^2^ < 0.01). For RTs, participants were overall faster in responding to incongruent trials on day 2, independent of the type of the preceding trial (main effect day: *F*_(1, 34)_ = 29.81, *p* < .001, η^2^ = 0.05), but we found no evidence for a Gratton effect on RTs (main effect preceding trial type *F* = 1.99, *p* = .167, η^2^ < 0.01) or a modulation of this effect across days (*F* = 3.07, *p* = .089, η^2^ < 0.01).

We did not observe significant behavioral adjustment after erroneous responses (“Laming effect”) in terms of accuracy, although there was a trend for improved performance following errors (main effect preceding accuracy: *F*_(1, 21)_ = 4.27, *p* = .051, η^2^_G_ = 0.32, main effect day: *F*_(1, 21)_ = 0.10, *p* = .754, η^2^_G_ < 0.01; preceding accuracy × day interaction: *F*_(1, 21)_ = 0.09, *p* = .770, η^2^_G_ < 0.01). For RTs, however, participants displayed significant post-error slowing (*F*_(1, 21)_ = 11.77, *p* = .003, η^2^_G_ = 0.05). While participants generally responded faster to incongruent trials on day 2 (*F*_(1, 21)_ = 17.54, *p* < .001, η^2^_G_ = 0.06), the Rabbitt effect on RTs was not modulated by day (preceding accuracy × day interaction: *F*_(1, 21)_ = 0.95, *p* = .341, η^2^_G_ < 0.01).

With respect to split-half reliability, estimates for accuracy ranged from acceptable to good in congruent (day 1: *r_split-half_* = .71; day 2: *r_split-half_* = .82) and incongruent trials (day 1: *r_split-half_* = .88; day 2: *r_split-half_* = .83) on both days. For RTs, split-half reliability was overall excellent for both congruent (day 1: *r_split-half_* = .98; day 2: *r_split-half_* = .99) and incongruent trials (day 1: *r_split-half_* = .99; day 2: *r_split-half_* = .98). Similarly, estimates for the flanker interference effect suggested acceptable to excellent split-half reliability for accuracy (day 1: *r_split-half_* = .88; day 2: *r_split-half_* = .74) and good to excellent split-half reliability for RTs (day 1: *r_split-half_* = .91; day 2: *r_split-half_* = .86). In contrast, split-half reliability was worse for the Gratton effect for both accuracy (day 1: *r_split-half_* = .63; day 2: *r_split-half_* = .01) and RTs (day 1: *r_split-half_* = .12; day 2: *r_split-half_* = .03). Estimates for the Laming/Rabbitt effects were numerically larger but still suggested low split-half reliability overall for both accuracy (day 1: *r_split-half_* = .19; day 2: *r_split-half_* = .29) and RTs (day 1: *r_split-half_* = .77; day 2: *r_split-half_* = .38).

As summarized in Table 1 and Figure 3, test-retest reliability estimates varied substantially between behavioral indices. Accuracy in congruent trials displayed lower reliability (*r* = ICC_3_ = .60, ICC_2_ = .61) than in incongruent trials (*r* = .89, ICC_3_ = ICC_2_ = .88), presumably because participants performed close to ceiling in congruent trials so that very small changes in performance between testing days may result in larger changes in the rank order. Both trial types showed low CV values (congruent: 0.83%, incongruent: 2.53%), indicating that participants’ absolute accuracy scores were very similar across days. RTs in both trial types showed good to excellent test-retest reliability (congruent: *r* = .93, ICC_3_ = .90, ICC_2_ = .83; incongruent: *r* = .89, ICC_3_ = .87, ICC_2_ = .76), with absolute RTs varying little between days (CV_congruent_ = 4.14%; CV_incongruent_ = 4.19%). Flanker interference effects displayed generally good test-retest reliability for both accuracy (*r* = ICC_3_ = ICC_2_ = .82) and RTs (*r* = ICC_3_ = .83, ICC_2_ = .81). In line with the poor internal consistency of both the Gratton and Laming/Rabbitt effects, test-retest reliability estimates for these indices were equally low. Specifically, the Gratton effect displayed poor test-retest reliability (accuracy: *r* = −.25, ICC_3_ = −.26, ICC_2_ = −.25; RT: *r* = ICC_3_ = .35, ICC_2_ = .34), with scores showing variability of CV_accuracy_ = 62.78% and CV_RT_ = 67.59%, respectively. The Laming/Rabbitt effects displayed poor test-retest reliability as well (accuracy: *r* = ICC_3_ = ICC_2_ =.15; RT: *r* = .43, ICC_3_ = .36, ICC_2_ = .37). While absolute scores for accuracy showed a variability of CV = 43.60%, variability for RTs was considerably lower (CV = 4.04%).

**Figure 3.**
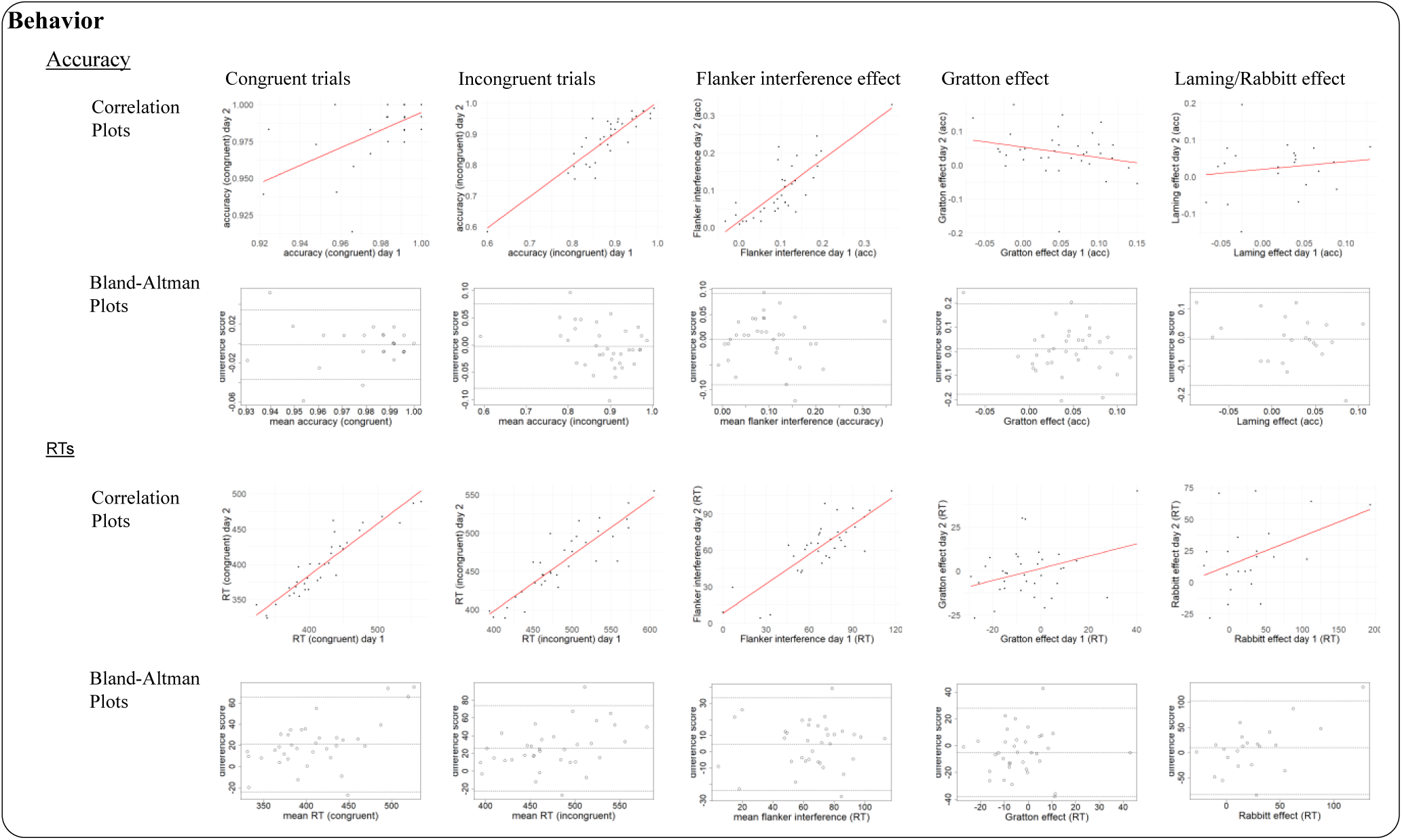
Pearson correlations and Bland-Altman plots for behavioral measures. Correlations were stronger for condition-specific variables (congruent and incongruent accuracy and RTs) than for difference scores. Similarly, the latter produced larger ranges for the 95% limits of agreement in the Bland-Altman plots than the former.

**Table 1.** Summary statistics, test-retest reliability, and repeatability estimates for behavioral markers in the Eriksen Flanker Task.

| | Average group performance<br>(mean ± SD) | | Test-retest reliability | | | Repeatability | |
| --- | --- | --- | --- | --- | --- | --- | --- |
| | Day 1 | Day 2 | $r$<br>[95% CI] | ICC<sub>3</sub><br>[95% CI] | ICC<sub>2</sub><br>[95% CI] | Mean CV<br>[range] in % | Mean difference<br>[95% limits of agreement] |
| Accuracy (in %) |  |  |  |  |  |  |  |
| Congruent trials | 98.37 ± 2.06 | 98.49 ± 2.04 | .60<br>[.34 – .78] | .60<br>[.34 – .78] | .61<br>[.35 – .78] | 0.83<br>[0.00 – 4.36] | -0.12<br>[-3.69 – 3.46] |
| Incongruent trials | 88.11 ± 7.43 | 88.28 ± 8.54 | .89<br>[.79 - .94] | .88<br>[.77 - .94] | .88<br>[.78 - .94] | 2.53<br>[0.09 – 8.47] | -0.17<br>[-7.88 – 7.54] |
| Flanker Interference<br>(incongruent – congruent) | 10.26 ± 7.59 | 10.21 ± 7.71 | .82<br>[.67 - .91] | .82<br>[.67 - .90] | .82<br>[.67 - .91] | / | 0.05<br>[-9.02 – 9.12] |
| Gratton effect<br>(post incongruent – post congruent) | 4.83 ± 5.42 | 3.82 ± 6.62 | -.26<br>[-.54 - .08] | -.25<br>[-.54 - .09] | -.26<br>[-.55 - .09] | / | 1.01<br>[-17.74 – 19.77] |
| Laming/ effect<br>(post error – post correct response) | 0.02 ± 0.05 | 0.02 ± 0.07 | .15<br>[-.29 - .54] | .15<br>[-.30 - .54] | .15<br>[-.28 - .53] | / | -0.52<br>[-16.80 – 15.75] |
| RTs (in ms) |  |  |  |  |  |  |  |
| Congruent trials | 420.34 ± 57.33 | 399.28 ± 45.06 | .93<br>[.86 - .96] | .90<br>[.82 - .95] | .83<br>[.36 - .94] | 4.14<br>[0.06 – 10.5] | 21.06<br>[-23.65 – 65.77] |
| Incongruent trials | 487.74 ± 52.76 | 461.91 ± 43.41 | .89<br>[.79 - .94] | .87<br>[.76 - .93] | .76<br>[.14 - .92] | 4.19<br>[0.30 – 13.16] | 25.83<br>[-22.29 – 73.94] |
| Flanker Interference<br>(incongruent – congruent) | 67.40 ± 24.86 | 62.63 ± 24.34 | .83<br>[.68 - .91] | .83<br>[.68 - .91] | .81<br>[.66 - .90] | / | 4.77<br>[-23.72 – 33.25] |
| Gratton effect (post congruent – post incongruent) | -5.38 ± 14.85 | -0.41 ± 14.65 | .35<br>[.02 - .61] | .35<br>[.03 - .61] | .34<br>[.03 - .60] | / | -4.98<br>[-37.91 – 27.95] |
| Rabbit effect (post error – post correct response) | 30.48 ± 52.87 | 20.52 ± 28.02 | .43<br>[.01 - .72] | .36<br>[-.07 - .67] | .36<br>[-.06 - .67] | / | 9.97<br>[-83.99 – 103.93] |

\* $r$ = Pearson correlation, ICC = Intraclass Correlation Coefficient, CV = Coefficient of Variation. Differences for repeatability are calculated as day 1 – day2.

With the exception of congruent and incongruent RTs (see ANOVA results), across-session differences centered close to zero, indicating the absence of systematic measurement differences between testing days. Moreover, only very few difference scores fell outside the 95% limits of agreement. However, the range of values covered by these limits was much larger for flanker interference, Gratton, and Laming/Rabbitt effects than for condition-specific accuracy and RTs, indicating higher heterogeneity in the sample for these differences scores.

### 3.2 EEG measures

#### 3.2.1 N200

##### 3.2.1.1 ERP

Contrary to our expectations, we did not observe a modulation of N200 magnitude by trial type (*F*_(1, 33)_ = 0.01, *p* = .958, η^2^_G_ < 0.01). Participants exhibited a larger (i.e., more negative) N200 amplitude on day 2 compared to day 1 (*F*_(1, 33)_ = 22.26, *p* < .001, η^2^_G_ = 0.03), but there was no trial type × day interaction (*F*_(1, 33)_ = 0.02, *p* = .891, η^2^_G_ < 0.01; see Figure 4A,B). Split-half reliability estimates for the N200 wave were excellent for both congruent (day 1: *r_split-half_* = .98; day 2: *r_split-half_* = .98) and incongruent trials (day 1: *r_split-half_* = .96; day 2: *r_split-half_* > .99).

**Figure 4.**
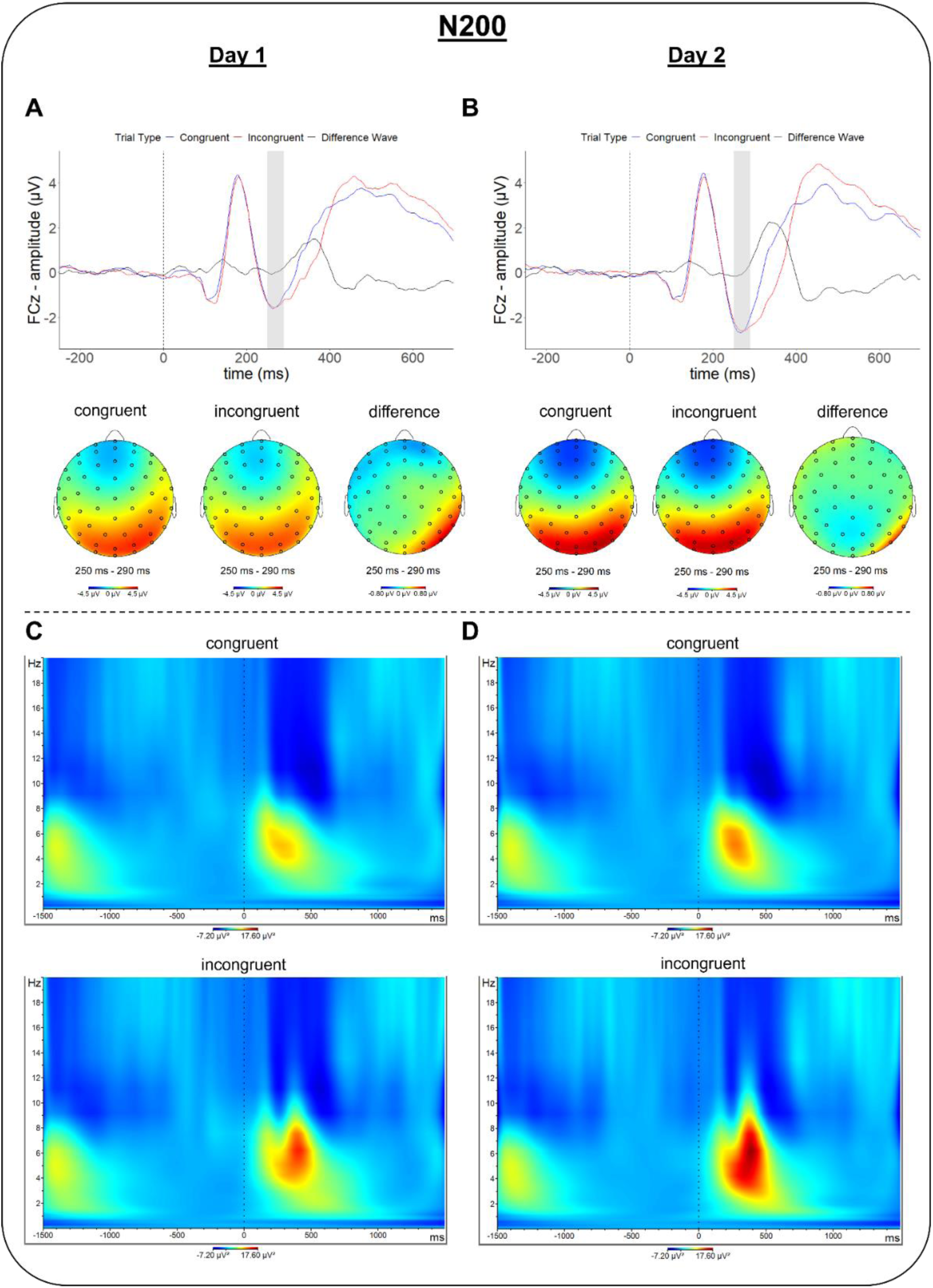
Target-locked ERP waves, scalp topographies, and spectral power in the N200 time-window (250 – 290ms after flanker onset). ERPs were measured at electrode channel 2 (∼FCz), depicting neural responses as well as scalp topographies following congruent and incongruent trials on day 1 (A) and day 2 (B). Difference scores were calculated as incongruent minus congruent conditions. Time-frequency plots depict spectral power in congruent and incongruent trials for day 1 (C) and day 2 (D). Grey overlays in ERP waveform plots (panels A and B) highlight the time-window used for analysis.

As seen in Table 2 and Figure 5, N200 amplitudes showed good to excellent test-retest reliability for both congruent (*r* = ICC_3_ = .93, ICC_2_ = .89) and incongruent trials (*r* = ICC_3_ = .91, ICC_2_ = .87). The limits of agreement for across-day difference scores were relatively large in both conditions. The N200 difference wave displayed overall poor test-retest reliability (*r* = ICC_3_ = ICC_2_ = .05) but displayed tighter limits of agreement.

**Figure 5.**
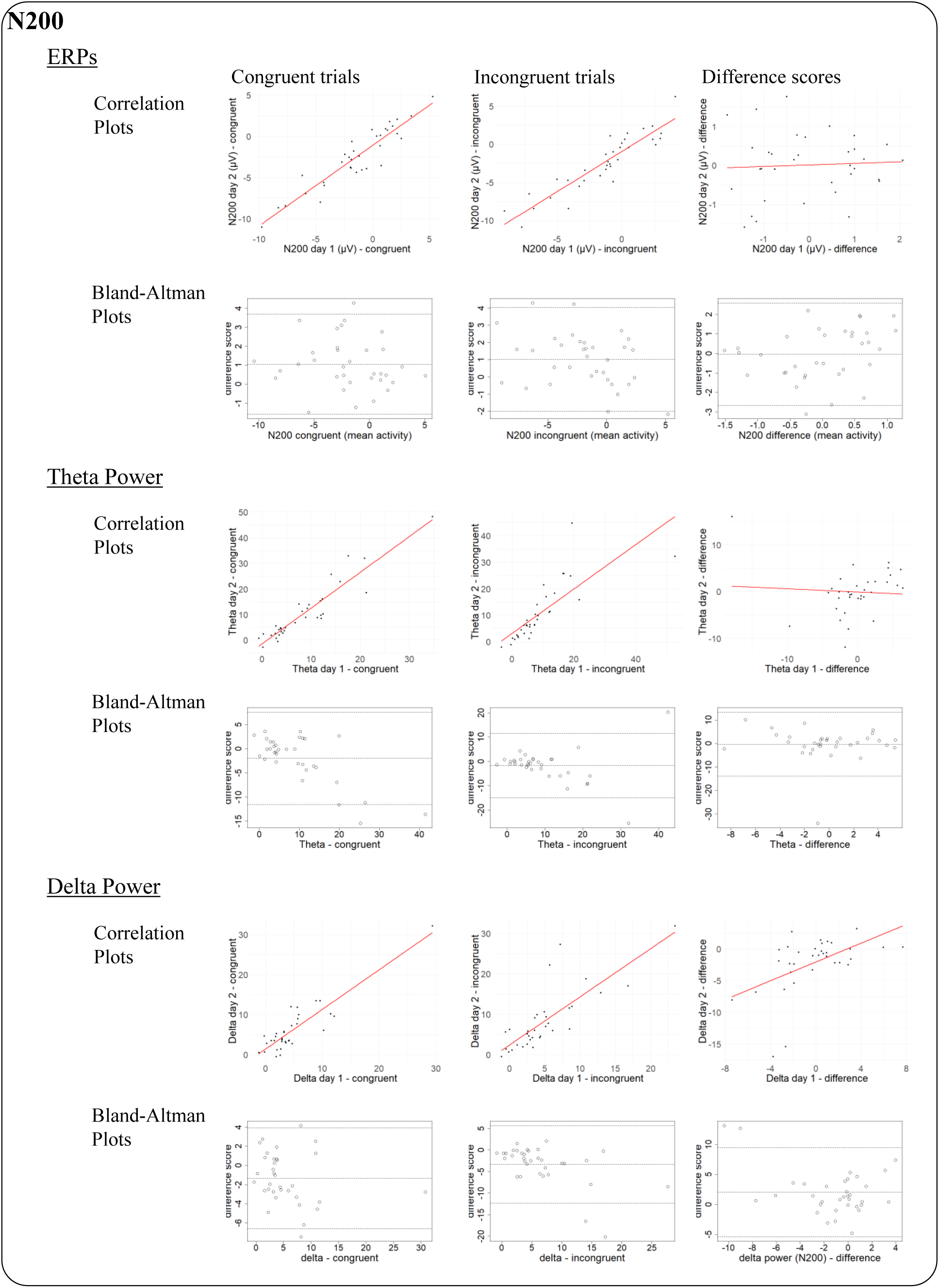
Pearson correlations and Bland-Altman plots for the N200 ERP component and associated spectral power. Correlations were stronger for condition-specific variables (congruent and incongruent trials) than for difference scores. Similarly, the latter produced larger ranges for the 95% limits of agreement in the Bland-Altman plots than the former. Across-task differences in delta power showed the strongest deviations from zero for both condition-specific values and between conditions. Given the lack of an expected congruency effect on the N200, reliability estimates may not generalize to other experimental tasks.

**Table 2.**
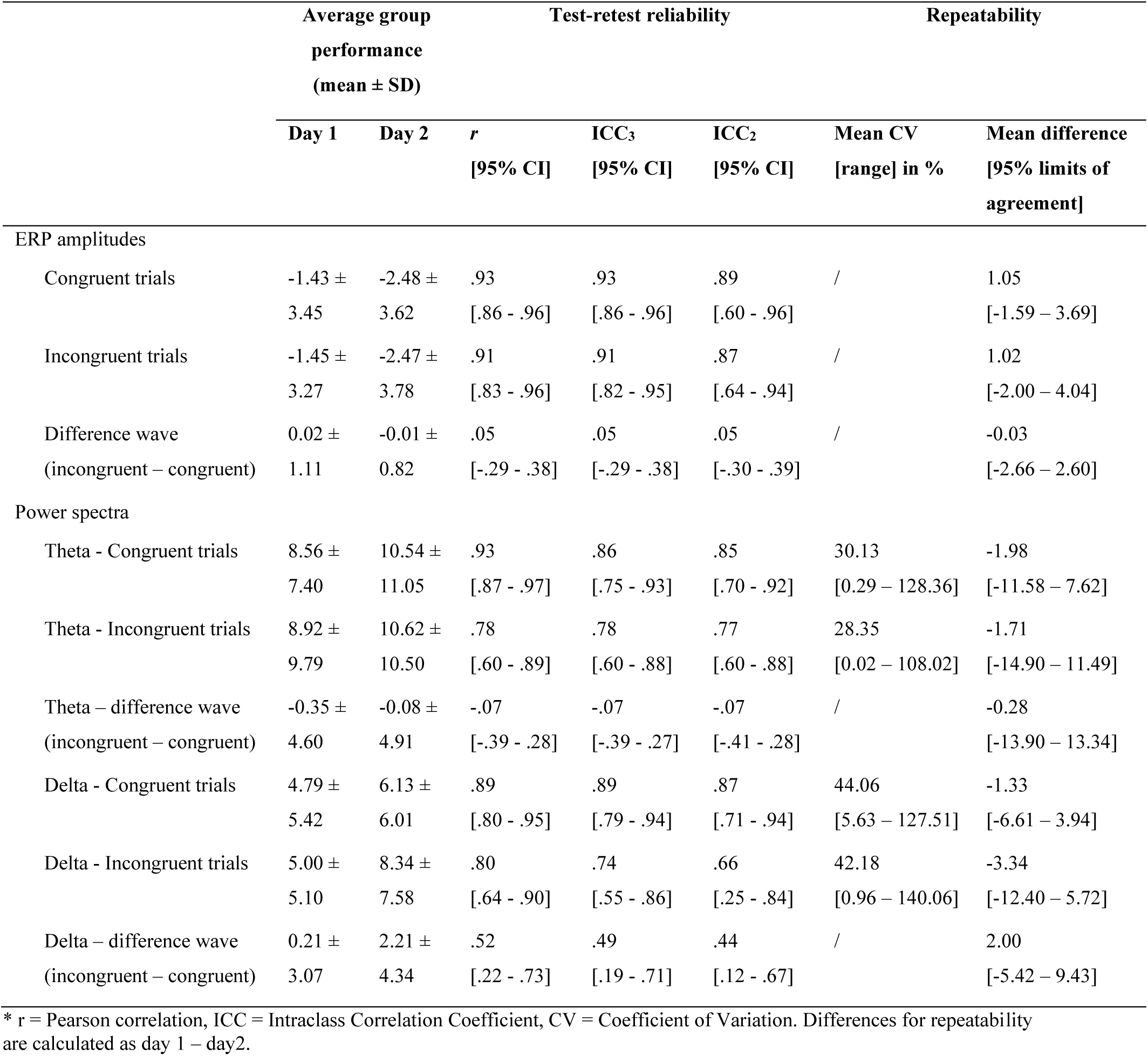
Summary statistics, test-retest reliability, and repeatability estimates for N200 markers in the Eriksen Flanker Task.

##### 3.2.1.2. Spectral power

Mirroring the ERP results, we found larger theta band power on day 2 compared to day 1 (*F*_(1, 33)_ = 5.12, *p* = .030, η^2^_G_ = 0.01) but did not observe the expected effect of trial type (*F*_(1, 33)_ = 0.15, *p* = .700, η^2^_G_ < 0.01) nor a trial type × day interaction (*F*_(1, 33)_ = 0.05, *p* = .818, η^2^_G_ < 0.01; see Figure 4C,D). For the delta band, we found that spectral power was larger for incongruent compared to congruent trials (*F*_(1, 33)_ = 4.73, *p* = .037, η^2^_G_ = 0.01) and larger on day 2 compared to day 1 (*F*_(1, 33)_ = 17.31, *p* < .001, η^2^_G_ = 0.04). Moreover, the difference in delta power between congruent and incongruent trials was larger on day 2 compared to day 1 (trial type × day interaction: *F*_(1, 33)_ = 9.51, *p* = .004, η^2^_G_ = 0.01).

Both theta and delta power exhibited good to excellent test-retest reliability (theta: *r* = .93, ICC_3_ = .86, ICC_2_ = .85; delta: *r* = ICC_3_ = .89, ICC_2_ = .87) in congruent trials, and acceptable to good test-retest reliability (theta: *r* = ICC_3_ =.78, ICC_2_ =.77; delta: *r* = .80, ICC_3_ =.74, ICC_2_ = .66) in incongruent trials. Absolute scores varied slightly less for theta power (congruent: CV = 30.13%; incongruent: CV = 28.35%) than for delta power (congruent: CV = 44.06%; incongruent: CV = 42.18%), which was mirrored in the range of the limits of agreement for these measures. Difference scores for the theta band showed overall poor test-retest reliability (*r* = ICC_3_ = ICC_2_ = −.07). Difference scores for the delta band showed higher, but still overall poor, estimates of test-retest reliability (*r* = .52, ICC_3_ = .49, ICC_2_ = .44).

### 3.3 Response-locked components

#### 3.3.1 ERN/CRN

##### 3.3.1.1 ERP

As expected, participants displayed a significantly larger (i.e., more negative) ERP amplitude after responding incorrectly than correctly (*F*_(1, 23)_ = 127.92, *p* < .001, η^2^ = 0.30; see Figure 6A,B). We did not observe a day (*F*_(1, 23)_ < 0.01, *p* = .986, η^2^ < 0.01) nor a response type × day interaction effect (*F*_(1, 23)_ = 0.05, *p* = .816, η^2^ < 0.01). Split-half reliability estimates for the ERN/CRN wave were excellent for correct (day 1: *r_split-half_* = .96; day 2: *r_split-half_* = .96) and acceptable to good for incorrect trials (day 1: *r_split- half_* = .84; day 2: *r_split-half_* = .75).

**Figure 6.**
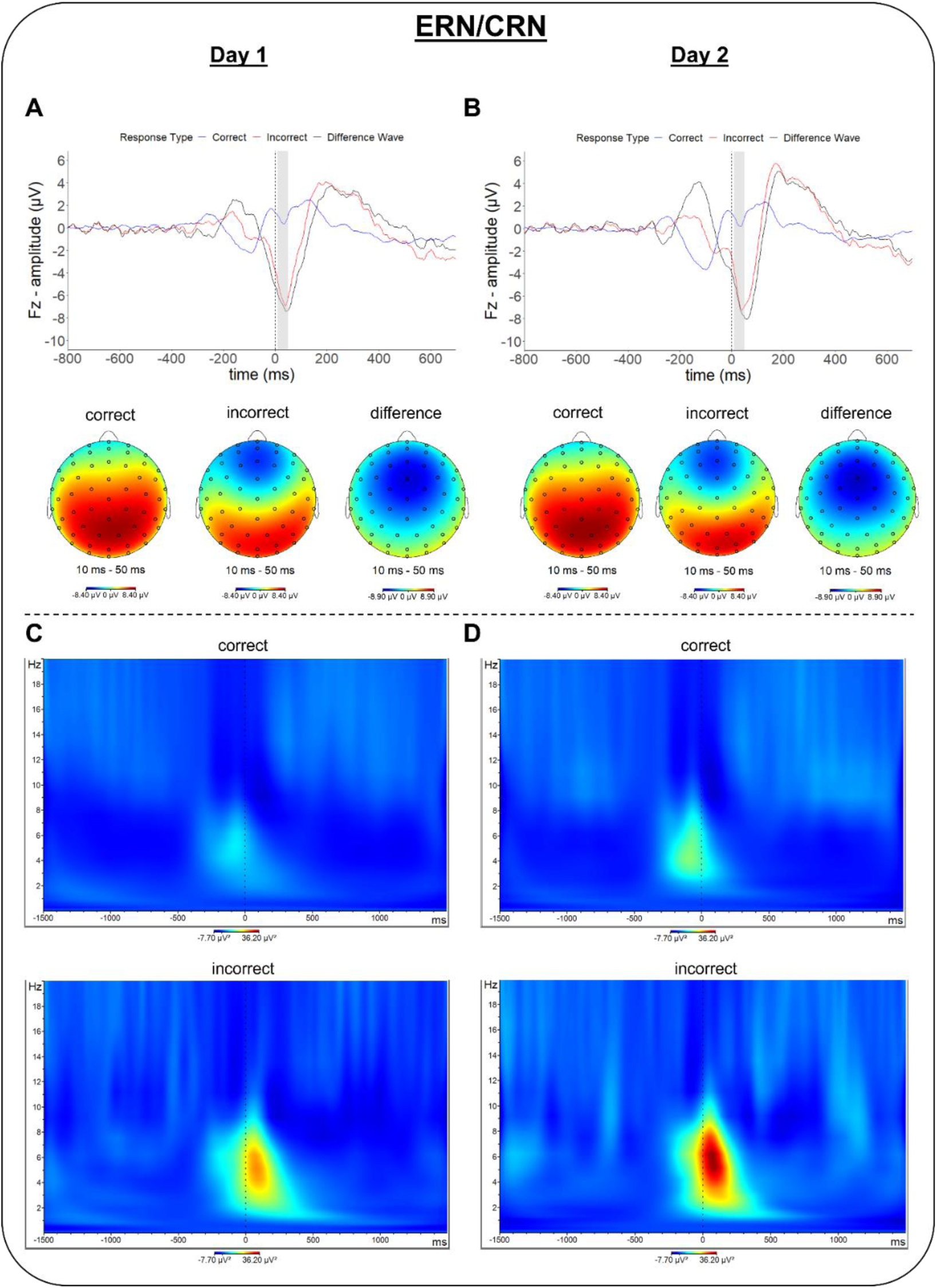
Response-locked ERP waves, scalp topographies, and spectral power in the ERN/CRN time-window (10 – 50ms after response). ERPs were measured at electrode channel 9 (∼Fz), depicting neural responses as well as scalp topographies following correct and i7ncorrect trials on day 1 (A) and day 2 (B). Difference scores were calculated as incorrect minus correct responses. Time-frequency plots depict spectral power in correct and incorrect trials for day 1 (C) and day 2 (D). Grey overlays in ERP waveform plots (panels A and B) highlight the time-window used for analysis.

As summarized in Table 3 and Figure 7, estimates for test-retest reliability were excellent for CRN amplitudes (*r* = ICC_3_ = ICC_2_ = .92) but overall poor for ERN amplitudes (*r* = ICC_3_ = .50, ICC_2_ = .51), presumably due a low amount of error trials across participants. The ERN/CRN difference wave displayed overall poor consistency and agreement (*r* = ICC_3_ = ICC_2_ = .01). Across-day differences centered around zero for both condition-specific and difference scores, however, ERN amplitudes displayed much more variability compared to CRN amplitudes and the difference wave.

**Figure 7.**
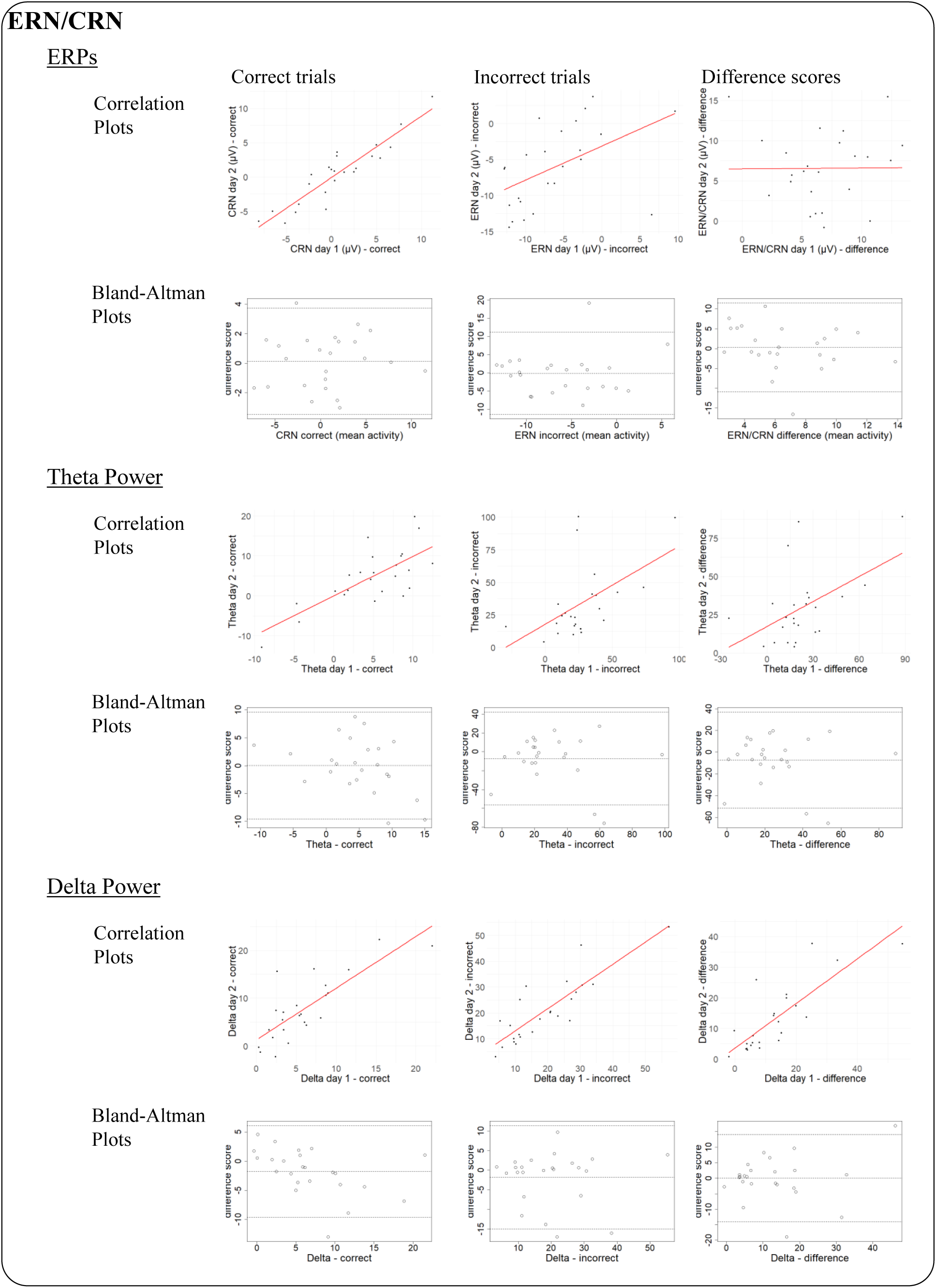
Pearson correlations and Bland-Altman plots for the CRN/ERN ERP component and associated spectral power. Correlations were strongest for correct responses compared to error responses or difference scores, except for the delta band, which showed overall good reliability. Similarly, error responses produced larger ranges for the 95% limits of agreement in the Bland-Altman plots compared to correct responses.

**Table 3.** Summary statistics, test-retest reliability, and repeatability estimates for ERN/CRN markers in the Eriksen Flanker Task.

| | Average group performance<br>(mean ± SD) | | Test-retest reliability | | | Repeatability | |
| --- | --- | --- | --- | --- | --- | --- | --- |
|  | Day 1 | Day 2 | *r*<br>[95% CI] | ICC<sub>3</sub><br>[95% CI] | ICC<sub>2</sub><br>[95% CI] | Mean CV<br>[range] in % | Mean difference<br>[95 % limits of agreement] |
| ERP amplitudes |  |  |  |  |  |  |  |
| Correct trials | 0.64 ± 4.60 | 0.49 ± 4.47 | .92<br>[.82 - .96] | .92<br>[.82 - .96] | .92<br>[.83 - .96] | / | 0.15<br>[-3.44 – 3.74] |
| Incorrect trials | -6.17 ± 5.89 | -6.05 ± 5.59 | .50<br>[.12 - .75] | .50<br>[.13 - .75] | .51<br>[.13 - .75] | / | -0.13<br>[-11.41 – 11.15] |
| Difference wave<br>(incorrect – correct) | 6.82 ± 3.56 | 6.54 ± 4.53 | .01<br>[-.40 - .41] | .01<br>[-.39 - .40] | .01<br>[-.41 - .41] | / | 0.28<br>[-10.97 – 11.52] |
| Power spectra |  |  |  |  |  |  |  |
| Theta – Correct trials | 4.78 ± 5.37 | 4.74 ± 7.19 | .73<br>[.47 - .88] | .70<br>[.42 - .86] | .71<br>[.43 - .86] | 48.82<br>[1.77 – 141.00] | 0.04<br>[-9.55 – 9.63] |
| Theta - Incorrect trials | 27.08 ± 24.60 | 34.14 ± 27.38 | .54<br>[.17 - .77] | .54<br>[.18 - .77] | .53<br>[.18 - .76] | 37.15<br>[2.16 – 88.30] | -7.06<br>[-56.21 – 42.08] |
| Theta – difference wave<br>(incorrect – correct) | 22.30 ± 22.00 | 29.40 ± 23.49 | .51<br>[.13 - .76] | .51<br>[.14 - .75] | .50<br>[.14 - .74] | / | -7.10<br>[-51.31 – 37.10] |
| Delta - Correct trials | 6.05 ± 4.96 | 7.82 ± 6.69 | .80<br>[.59 - .91] | .77<br>[.53 - .89] | .74<br>[.47 - .88] | 32.64<br>[0.93 – 104.27] | -1.77<br>[-9.65 – 6.11] |
| Delta – Incorrect trials | 19.08 ± 12.19 | 20.88 ± 12.31 | .85<br>[.68 - .93] | .85<br>[.68 - .93] | .85<br>[.68 - .93] | 17.01<br>[0.42 – 74.01] | -1.80<br>[-14.97 – 11.38] |
| Delta – difference wave<br>(incorrect – correct) | 13.03 ± 12.29 | 13.06 ± 11.02 | .81<br>[.61 - .92] | .81<br>[.61 - .91] | .82<br>[.62 - .92] | / | -0.03<br>[-14.12 – 14.06] |

\* *r* = Pearson correlation, ICC = Intraclass Correlation Coefficient, CV = Coefficient of Variation. Differences for repeatability are calculated as day 1 – day 2.

##### 3.3.1.2 Spectral power

Similar to the ERPs, we found that spectral power in the theta band was, as expected, significantly higher for error compared to correct responses (*F*_(1, 23)_ = 41.04, *p* < .001, η^2^_G_ = 0.33; see Figure 6C,D). This effect was not modulated by day (main effect day: *F*_(1, 23)_ = 1.48, *p* = .235, η^2^_G_ = 0.01; response type × day interaction: *F*_(1, 23)_ = 2.38, *p* = .137, η^2^_G_ = 0.01). In the delta band, spectral power was larger for error compared to correct responses as well, in line with our predictions (*F*_(1, 23)_ = 33.13, *p* < .001, η^2^_G_ = 0.32). Although delta power was higher on day 2 (*F*_(1, 23)_ = 4.31, *p* = .049, η^2^_G_ = 0.01), we found no significant response type × day interaction (*F*_(1, 23)_ < 0.01, *p* = .985, η^2^_G_ < 0.01).

While both theta and delta band power exhibited acceptable test-retest reliability (theta: *r* = .73, ICC_3_ = .70, ICC_2_ = .71; delta: *r* = .80, ICC_3_ = .77, ICC_2_ = .74), these estimates were poor for theta band power in error trials (*r* = ICC_3_ = .54, ICC_2_ = .53). In contrast, delta power in error trials exhibited overall good test-retest reliability (*r* = ICC_3_ = ICC_2_ = .85). Similarly, test-retest reliability measures for difference scores between correct and error responses were poor for theta (*r* = ICC_3_ = .51, ICC_2_ = .50) but good for delta power (*r* = ICC_3_ = .81, ICC_2_ = .82). On average, absolute values varied slightly more for theta power (correct: CV = 48.82%; incorrect: CV = 37.15%) compared to delta power (correct: CV = 32.64%; incorrect: CV = 17.01%). As for the ERPs, theta and delta power in response to errors produced wider limits of agreement than those following correct responses.

##### 3.3.2 Pe

###### 3.3.2.1 ERP

In line with previous research, we found that errors led to a significantly larger (i.e., more positive) ERP amplitude compared to correct responses (*F*_(1, 23)_ = 11.66, *p* = .002, η^2^ = 0.16; see Figure 8A,B). We did not observe a main effect of day (*F*_(1, 23)_ = 0.65, *p* = .428, η^2^ < 0.01) or a response type × day interaction (*F*_(1, 23)_ = 2.80, *p* = .108, η^2^ < 0.01). Split-half reliability estimates for the Pe wave were excellent for correct (day 1: *r_split-half_* = .94; day 2: *r_split-half_* = .93) and good to excellent for incorrect trials (day 1: *r_split-half_* = .89; day 2: *r_split-half_* = .94).

**Figure 8.**
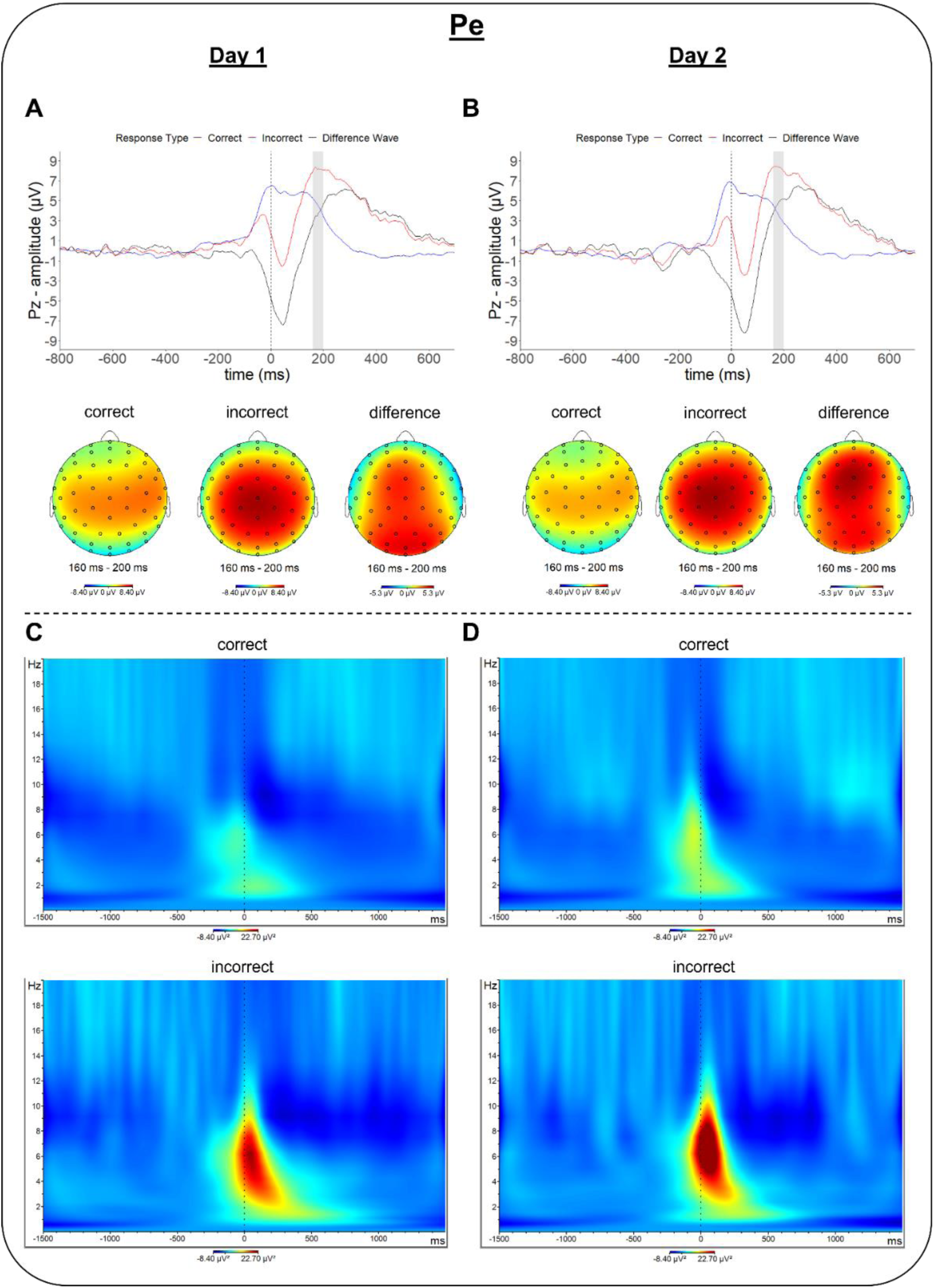
Response-locked ERP waves, scalp topographies, and spectral power in the Pe time-window (160 – 200ms after response). ERPs were measured at electrode channel 1 (∼Cz), depicting neural responses as well as scalp topographies following correct and incorrect trials on day 1 (A) and day 2 (B). Difference scores were calculated as incorrect minus correct responses. Time-frequency plots depict spectral power in correct and incorrect trials for day 1 (C) and day 2 (D). Grey overlays in ERP waveform plots (panels A and B) highlight the time-window used for analysis.

As summarized in Table 4 and Figure 9, estimates for test-retest reliability were acceptable to good for Pe amplitudes in correct trials (*r* = .81, ICC_3_ = .80, ICC_2_ = .78) and good for incorrect trials (*r* = ICC_3_ = .87, ICC_2_ = .88). The Pe difference wave displayed good consistency and agreement (*r* = ICC_3_ = ICC_2_ = .89). Bland-Altman plots showed that across-day differences were centered near zero and limits of agreement were relatively tight across all conditions.

**Figure 9.**
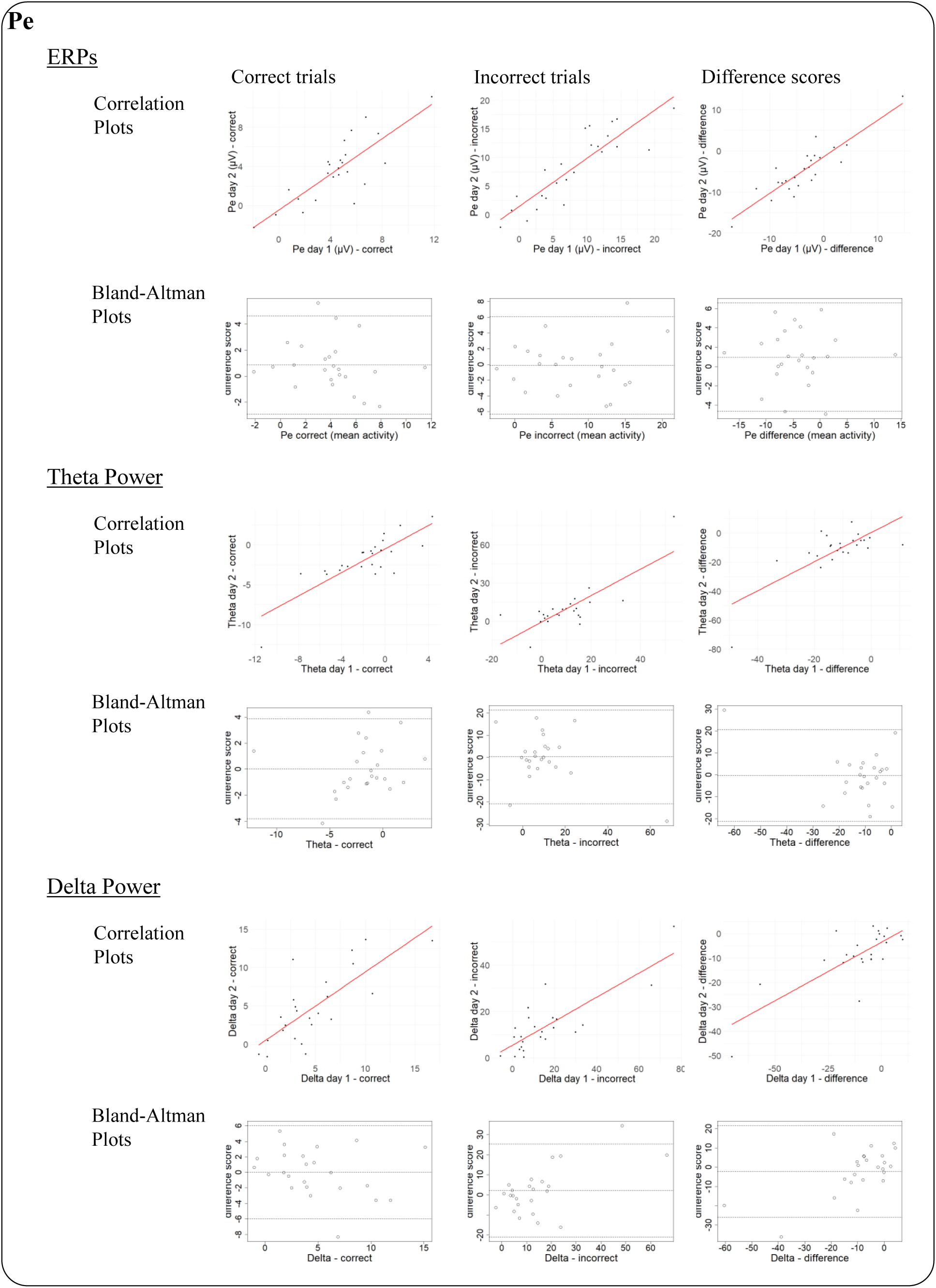
Pearson correlations and Bland-Altman plots for the Pe ERP component and associated spectral power. Compared to other ERPs, correlations for Pe amplitudes were stronger and more consistent across condition-specific variables (correct and incorrect responses) and difference scores. The 95% limits of agreement for ERPs in the Bland-Altman plots were relatively tight across conditions but were wider for error responses and difference scores compared to correct responses in both frequency bands.

**Table 4.** Summary statistics, test-retest reliability, and repeatability estimates for Pe markers in the Eriksen Flanker Task.

| | Average group performance<br>(mean ± SD) | | Test-retest reliability | | | Repeatability | |
| --- | --- | --- | --- | --- | --- | --- | --- |
|  | Day 1 | Day 2 | *r*<br>[95% CI] | ICC<sub>3</sub><br>[95% CI] | ICC<sub>2</sub><br>[95% CI] | Mean CV<br>[range] in % | Mean difference<br>[95% limits of agreement] |
| ERP amplitudes |  |  |  |  |  |  |  |
| Correct trials | 4.48 ± 2.87 | 3.62 ± 3.25 | .81<br>[.60 - .91] | .80<br>[.60 - .91] | .78<br>[.54 - .90] | / | 0.85<br>[-2.93 – 4.63] |
| Incorrect trials | 8.18 ± 6.41 | 8.31 ± 6.17 | .87<br>[.72 - .94] | .87<br>[.73 - .94] | .88<br>[.74 - .95] | / | -0.13<br>[-6.33 – 6.07] |
| Difference wave<br>(correct – incorrect) | -3.71 ± 6.20 | -4.69 ± 6.19 | .89<br>[.76 - .95] | .89<br>[.77 - .95] | .89<br>[.75 - .95] | / | 0.98<br>[-4.65 – 6.61] |
| Power spectra |  |  |  |  |  |  |  |
| Theta – Correct trials | -1.84 ± 3.42 | -1.87 ± 3.06 | .82<br>[.63 - .92] | .82<br>[.63 - .92] | .82<br>[.64 - .92] | 49.85<br>[6.23 – 132.27] | 0.04<br>[-3.80 – 3.87] |
| Theta - Incorrect trials | 10.13 ± 13.50 | 9.77 ± 17.67 | .79<br>[.57 - .91] | .77<br>[.53 - .89] | .77<br>[.54 - .90] | 47.99<br>[0.68 – 120.03] | 0.35<br>[-20.71 – 21.42] |
| Theta – difference wave<br>(correct – incorrect) | -11.96 ± 11.79 | -11.64 ± 15.87 | .74<br>[.49 - .88] | .71<br>[.44 - .86] | .72<br>[.45 - .87] | / | -0.32<br>[-21.09 – 20.45] |
| Delta - Correct trials | 4.80 ± 3.98 | 4.78 ± 4.66 | .76<br>[.51 - .89] | .75<br>[.50 - .88] | .76<br>[.51 - .89] | 43.29<br>[0.19 – 140.10] | 0.02<br>[-5.99 – 6.02] |
| Delta – Incorrect trials | 15.64 ± 19.54 | 13.47 ± 12.42 | .82<br>[.62 - .92] | .74<br>[.49 - .88] | .74<br>[.49 - .88] | 48.47<br>[4.17 – 124.33] | 2.17<br>[-20.97 – 25.31] |
| Delta – difference wave<br>(correct – incorrect) | -10.84 ± 18.66 | -8.68 ± 11.53 | .78<br>[.54 - .90] | .70<br>[.41 - .86] | .70<br>[.42 - .86] | / | -2.15<br>[-25.88 – 21.58] |

\* *r* = Pearson correlation, ICC = Intraclass Correlation Coefficient, CV = Coefficient of Variation. Differences for repeatability are calculated as day 1 – day 2.

###### 3.3.2.2 Spectral power

For the theta band, we found the expected significant main effect of response type (*F*_(1, 23)_ = 19.98, *p* < .001, η^2^_G_ = 0.22), due to greater spectral power in error compared to correct trials (see Figure 6C,D). No other effect was significant (main effect day: *F*_(1, 23)_ = 0.03, *p* = .866, η^2^_G_ < 0.01; response type × day interaction: *F*_(1, 23)_ = 0.02, *p* = .885, η^2^_G_ < 0.01). These findings were mirrored in the delta band. As expected, we observed greater power in error compared to correct trials (*F*_(1, 23)_ = 11.21, *p* = .003, η^2^_G_ = 0.15) but no significant effects involving day (main effect day: *F*_(1, 23)_ = 0.76, *p* = .392, η^2^_G_ < 0.01; response type × day interaction: *F*_(1, 23)_ = 0.76, *p* = .393, η^2^_G_ < 0.01).

Test-retest reliability measures of theta band activity were good for correct trials (*r* = ICC_3_ = ICC_2_ = .82) and acceptable for both incorrect trials (*r* = .79, ICC_3_ = ICC_2_ = .79) and the difference scores (*r* = .74, ICC_3_ = .71, ICC_2_ = .72). For delta power, test-retest reliability measures were acceptable to good for incorrect trials (*r* = .82, ICC_3_ = ICC_2_ = .74) and acceptable for both correct trials (*r* = .76, ICC_3_ = .75, ICC_2_ = .76) and difference scores (*r* = .78, ICC_3_ = .70, ICC_2_ = .70). On average, absolute values for theta (correct: CV = 49.85%; incorrect: CV = 47.99%) and delta power (correct: CV = 43.29%; incorrect: CV = 48.47%) varied to a similar degree. Across-day differences were centered near zero for all conditions in both frequency bands, however, error responses produced wider limits of agreement than correct responses.

## 4 Discussion

In this study, we investigated behavioral and electrophysiological correlates of interference control and performance monitoring in the classic Eriksen Flanker Task and evaluated their test-retest reliability and repeatability across 48 hours among psychiatrically healthy individuals.

### 4.1 Test-retest reliability

Our findings suggest that individual cognitive control metrics varied considerably, ranging from excellent to unacceptable (see Figure 10). In terms of test-retest reliability (Pearson’s *r*, ICC_3_ and ICC_2_), trial- or response-type specific indices displayed higher consistency than indices based on difference scores, which is in line with previous reports and usually explained by the fact that subtracting test scores increases the relative proportion of error variance to total score variance (Paap & Sawi, 2016).

**Figure 10.**
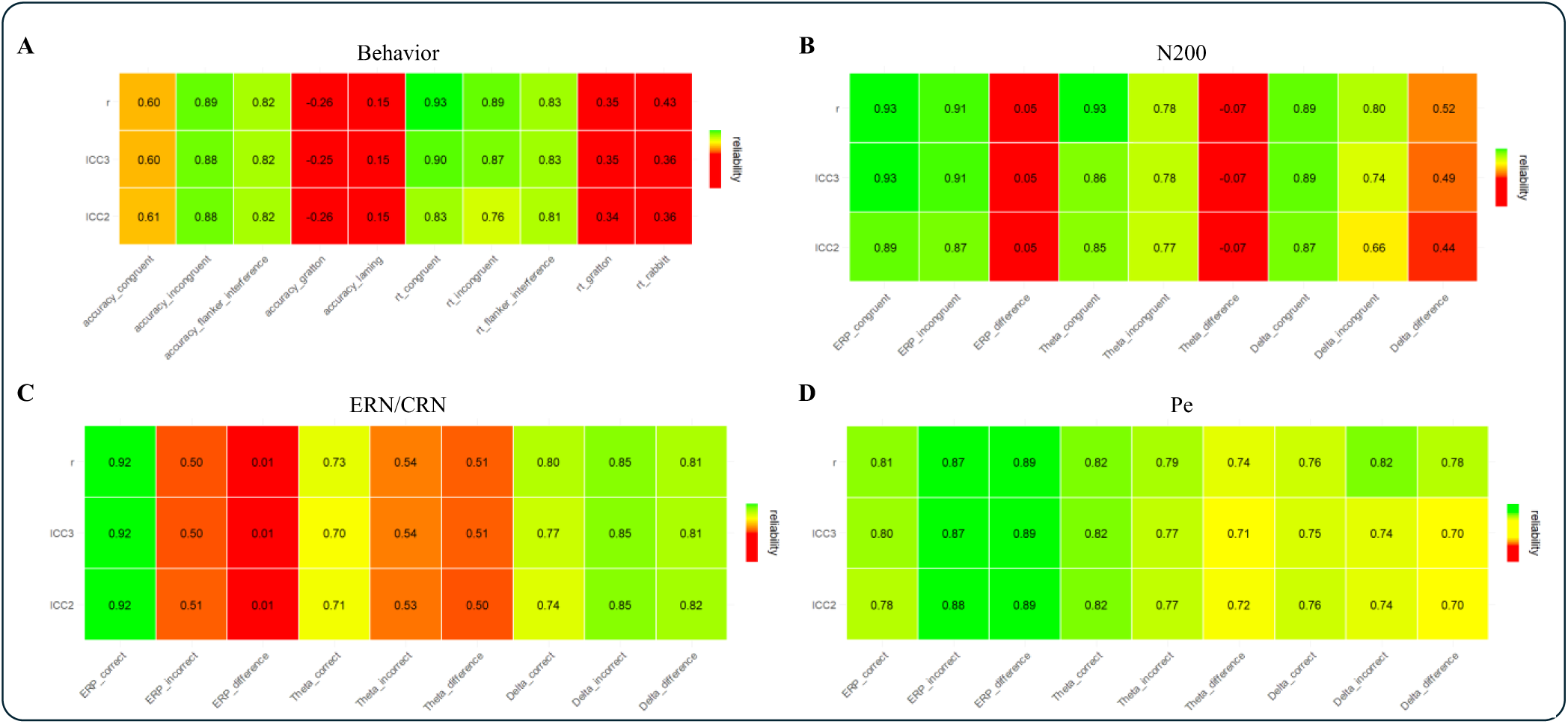
Heatmap of Test-Retest Reliability Estimates. For most behavioral and EEG-derived measures, condition-specific metrics showed higher reliability compared to differences scores. Overall, the Pe displayed the highest test-retest estimates. Colors indicate the psychometric quality of the coefficients from red (unacceptable) to green (good-to-excellent).

Behaviorally, RTs in both trial types and accuracy in incongruent trials were highly reliable, whereas accuracy in congruent trials was not, presumably due to the limited range of scores in these trials given that participants performed at ceiling. Flanker interference effects in both accuracy and RTs displayed good test-retest reliability, but it was poor for Gratton and Laming/Rabbitt effects. These findings align with the relatively few previous reports on the test-retest reliability of flanker performance measures in that RTs are commonly more reliable than accuracy scores (Lin et al., 2020; Paap & Sawi, 2016; Sanders et al., 2019; Wöstmann et al., 2013). Further, our estimates for accuracy and RTs in congruent and incongruent trials as well as the flanker interference in both outcomes suggested stronger test-retest reliability compared to earlier studies, which may be due to the comparatively short test-retest interval in our study. Test-retest estimates for Gratton and Laming/Rabbitt effects have not been frequently reported in the literature. However, based on our results, these indices seem generally inconsistent and may thus represent inferior choices for assessing individual differences or treatment efficacy related to cognitive control.

Similar to the behavioral findings, we observed that, except for the ERN, ERP magnitudes and spectral power in theta and delta frequency bands showed acceptable to excellent reliability when investigated within the same trial or response type and overall lower reliability when investigated as difference scores, which is in line with prior reports (Ip et al., 2018; Morand-Beaulieu et al., 2021; Olvet & Hajcak, 2009; Paap & Sawi, 2016). In contrast to both N200 and ERN/CRN-related measures, for both of which the ERP difference waves were almost completely inconsistent across days, Pe-related measures displayed generally high reliability. These findings are at odds with previous work investigating test-retest reliability in these components, which suggested moderate to good reliability for the difference waves of all of these components (Larson et al., 2010; Lin et al., 2020; Morand-Beaulieu et al., 2021; Olvet & Hajcak, 2009; Riesel et al., 2013; Weinberg & Hajcak, 2011). With respect to the N200, this discrepancy can be explained by the fact that we did not observe the expected modulation of ERP amplitudes and theta power by congruency reported in other studies (Kanske & Kotz, 2010; Kopp et al., 1996) (see below), which diminished the likelihood of these difference scores to be reliable across days. With respect to the ERN/CRN, the surprisingly low reliability of the ERP difference wave is likely due to the poor test-retest reliability of the condition-specific ERPs we observed for error responses, which, in turn, may be caused by the generally low number of incorrect trials in our version of the flanker task. Thus, a more difficult task might be needed to improve psychometrics of the ERN/CRN. Interestingly, our findings suggest that test-retest reliability of spectral power in the theta and delta frequencies closely aligns with estimates obtained for the ERPs in the corresponding time windows. Overall, our findings highlight the potential use of the Pe as a reliable marker of cognitive control in addition to the often-recommended ERN.

### 4.2 Repeatability

In addition to test-retest reliability, we were also interested in repeatability metrics to quantify the degree of intraindividual variability of cognitive control indices that provide important information about how much of a difference one can expect in absolute scores when repeatedly testing the same participants in the same task (Bartlett & Frost, 2008; Duda et al., 2021; Hopkins, 2000; Shechtman, 2013; Vaz et al., 2013). Our analysis suggests that for most of the measures considered here, exact scores will differ substantially across repeated assessments. In fact, only the basic, condition-specific behavioral indices (i.e., accuracy and RTs in either congruent or incongruent trials) fall below 5%, indicating highly similar scores in these outcomes across participants and days. One EEG-derived score displayed CVs between 10 – 20% (delta power following incorrect responses in the ERN time-window, CV = 17.01%), while all other outcomes produced CVs between 28.35 – 49.85%, which falls above thresholds previously suggested in studies investigating CVs specifically in medical imaging (Aronhime et al., 2014; Duda et al., 2021). It should be noted, however, that these values represent averages calculated across participants and that we observed considerable interindividual heterogeneity in CVs for all measures, often ranging from variability estimates of < 1% to > 100%, the latter indicating that, for a given person, the variability between testing days (i.e., the standard deviation) is larger than the mean score across days. Visual inspection of the Bland-Altman plots demonstrates that, while overall variability was large for the majority of measures, individual difference scores between day 1 and day 2 fell mostly within the 95% limits of agreement (i.e., ± 1.96 SD around the individual’s average score), indicating generally acceptable agreement of measurements across days (Altman & Bland, 1983). Further, for the majority of measures, the average difference score was close to zero, indicating that there was no systematic measurement bias between testing days. Notable exceptions to this observation are RTs in congruent and incongruent trials, which reflect the speeding of responses in both conditions on day 2 compared to day 1 and N200 magnitudes for congruent and incongruent trials, reflecting the larger amplitudes on day 2 compared to day 1.

While a certain degree of variability might be expected when investigating (neuro)psychological constructs, it may be challenging to assess whether changes in behavioral performance or EEG-related measures across time for a given individual, for example, following a treatment or experimental manipulation, can truly be attributed to these interventions or whether they represent natural fluctuations in these measures. To differentiate between these possibilities, it might thus be advisable to collect multiple baseline measurements to establish the individual variability in the construct of interest before administering and evaluating the efficacy of any intervention.

### 4.3 Behavioral and electrophysiological correlates of cognitive control

Generally, we replicated the majority of the expected behavioral and EEG-related effects commonly reported for flanker tasks. Specifically, on both days, participants displayed pronounced flanker interference effects for both accuracy and RTs, post-conflict adjustment (Gratton effects; accuracy only), and post-error adjustments (Laming/Rabbitt effects; significant for RTs only but descriptively present for accuracy as well). Similarly, the flanker task successfully and robustly elicited ERPs associated with error awareness and processing (i.e., more negative amplitudes in the ERN/CRN and more positive amplitudes in the Pe following incorrect compared to correct responses), as well as increased theta and delta power in error trials during both of these time windows (Eriksen, 1995; Olvet & Hajcak, 2009; Sandre & Weinberg, 2019). Notably, however, and in contrast to our hypothesis, we did not observe a modulation of N200 amplitude, nor theta power in the same time-window, by flanker congruency. This was somewhat surprising given the number of previous findings indicating a robust effect usually expressed as a substantially more negative deflection of this ERP in incongruent compared to congruent trial (Kanske & Kotz, 2010; Kopp et al., 1996)s. However, there is evidence suggesting that such trial type-related differences in N200 magnitude might not occur in flanker tasks in which congruent and incongruent trials appear with equal probability (Kałamała et al., 2018), as was the case in our study. Interestingly, we did observe significantly higher delta band power in incongruent compared to congruent trials, which may hint at a unique role of this frequency band in processing interference even in the absence of an observable N200 difference.

### 4.4 Limitations and conclusion

When interpreting our findings, one should keep in mind that for this study, participants completed their second testing session 48 hours after their first. On one hand, this comparatively short inter-test interval should have helped to increase the expected stability of the assessed behavioral and neural measures, as it is unlikely that participants’ capacity to exert cognitive control drastically changed within two days. On the other hand, completing the same task in quick succession could have resulted in training effects, which might have changed participants’ performance and, consequently, the relative or absolute outcomes in each of our measures. Indeed, participants demonstrated overall faster RTs (across both trial types) on day 2 compared to day 1 at similar accuracy levels. In line with this finding, average N200 amplitude and theta power (again across trial types) were larger on day 2, which may reflect an increase in task engagement, or effort, compared to day 1.

While we believe that our findings provide important insights into the psychometric properties of the flanker task and associated electrophysiological measures, we would like to note a few limitations. First, while our sample size may seem relatively modest, especially when taking into account further exclusions necessary for measures that depended on error responses, it matches samples from previous studies investigating test-retest reliability of behavioral and EEG-related outcomes as well as those commonly used in clinical or intervention studies (Button et al., 2013; Larson et al., 2010; Morand-Beaulieu et al., 2021; Olvet & Hajcak, 2009; Riesel et al., 2013; Weinberg & Hajcak, 2011). However, given that we tested only healthy adults in a predominantly White and mostly female sample, additional work is necessary to assess whether our findings generalize to other demographics or whether the test-retest reliability and repeatability of cognitive control indices vary more substantially across populations. Second, the fact that we, surprisingly, did not observe the expected N200 amplitude differences between congruent and incongruent trials makes it difficult to truly assess its reliability, given that reliability measures struggle when difference scores center around zero (Bartko, 1976). Finally, one of the central aims of this study was to assess not only the reliability of cognitive control indices, which assesses within-group variability, but also the degree to which scores of individual participants varied in absolute terms. In order to assess this, we calculated CVs, which are commonly used to measure repeatability (Berchtold, 2016; Duda et al., 2021; Hopkins, 2000; Shechtman, 2013; Vaz et al., 2013). However, given that CVs require measurements to lie on ratio scales, they may be somewhat limited in their use for many indices of cognitive processes that are often assessed by difference scores. However, we believe that quantifying this variability in absolute values is a useful metric to form expectations about how large changes caused, for example, by any kind of intervention would have to be to be considered meaningful for a given individual. Bland-Altman plots and the associated statistics (i.e., mean-difference score centered around zero, 95% limits of agreement) may provide additional information about the repeatability of a given measurement, as they can help to assess the expected range of variability in individual scores when participants are tested repeatedly and to estimate whether the absolute values of these scores change between repeated measures, even if the rank-order in the sample does not. We thus argue that future studies aimed at investigating the stability or malleability of cognitive processes over time should include these statistics in addition to the more traditional indices of test-retest reliability.

In summary, our findings suggest that most behavioral and electrophysiological measures of cognitive control in the classic Eriksen Flanker Task display satisfactory to excellent test-retest reliability across a 48-hour window, with condition-specific measures being generally more reliable than difference scores, especially those based on low trial counts. However, behavioral measures of post-conflict and post-error adjustment (i.e., Gratton and Laming/Rabbitt effects) were highly unreliable, as were difference scores for ERN and N200 amplitude as well as the underlying theta power, the latter of which resulted from a general lack of congruency-related differences in N200 amplitude. Further, by calculating CVs and Bland-Altman statistics, we found that absolute scores in all but the most basic behavioral indices varied considerably across time within participants and that the degree of this variability itself differed widely between participants. Thus, the results presented here not only provide novel information about the stability of cognitive and physiological markers of cognitive control over time but may also have implications for the application and evaluation of clinical interventions by helping to highlight and potentially to avoid obstacles in assessing the efficacy of novel treatment options in individuals. Most importantly, we suggest that researchers interested in tracking changes over time in clinically relevant domains and constructs, such as those defined in the RDoC framework (Insel, 2014), should be concerned not only with the reliability of their measure of interest, but also establish, on an individual level, how variable repeated measurements may be within single individuals and include these metrics in their reports.

## Conflict of interest

Over the past 3 years, Dr. Pizzagalli has received consulting fees from Boehringer Ingelheim, Compass Pathways, Engrail Therapeutics, Karla Therapeutics, Neumora Therapeutics (formerly BlackThorn Therapeutics), Neurocrine Biosciences, Neuroscience Software, Sage Therapeutics, Sama Therapeutics, and Takeda; he has received honoraria from the American Psychological Association, Psychonomic Society and Springer (for editorial work) and Alkermes; he has received research funding from the Bird Foundation, Brain and Behavior Research Foundation, Dana Foundation, DARPA, Millennium Pharmaceuticals, NIMH, and Wellcome Leap; he has received stock options from Compass Pathways, Engrail Therapeutics, Neumora Therapeutics, and Neuroscience Software. All views expressed are solely those of the authors. All other authors have no conflicts of interest or relevant disclosures.

## Funding

This work was supported by an investigator-initiated contract from Millennium Pharmaceuticals (awarded to DAP).

## Ethics statement

Study procedures were conducted in accordance with the Declaration of Helsinki and were approved by the MGB Healthcare Institutional Review Board

## Code/Data availability statement

### Author contributions

Conceptualization: DAP, RCM, BWB; Data curation: SRL, MB; Formal analysis: MB; Funding acquisition: DAP; Investigation: JNS, SME, ML, SEW; Methodology: DAP; Project administration: DAP; Resources: DAP, RCM; Supervision: DAP; Visualization: MB; Writing/original draft: MB, TL, DAP; Writing/review & editing: MB, JNS, SME, BWB, TL, ML, SEW, SRL, CM, SL, PB, RCM, DAP.

## Reference

Aldridge, V. K., Dovey, T. M., & Wade, A. (2017). Assessing test-retest reliability of psychological measures: Persistent methodological problems. European Psychologist, 22(4), 207.

Altman, D. G., & Bland, J. M. (1983). Measurement in medicine: The analysis of method comparison studies. Journal of the Royal Statistical Society Series D: The Statistician, 32(3), 307–317.

Aronhime, S., Calcagno, C., Jajamovich, G. H., Dyvorne, H. A., Robson, P., Dieterich, D., Isabel Fiel, M., Martel- Laferriere, V., Chatterji, M., Rusinek, H., & Taouli, B. (2014). DCE-MRI of the liver: Effect of linear and nonlinear conversions on hepatic perfusion quantification and reproducibility. Journal of Magnetic Resonance Imaging, 40(1), 90–98. 10.1002/jmri.24341

Bartko, J. J. (1966). The intraclass correlation coefficient as a measure of reliability. Psychological Reports, 19(1), 3–11.

Bartko, J. J. (1976). On various intraclass correlation reliability coefficients. Psychological Bulletin, 83(5), 762.

Bartlett, J., & Frost, C. (2008). Reliability, repeatability and reproducibility: Analysis of measurement errors in continuous variables. Ultrasound in Obstetrics & Gynecology, 31(4).

Berchtold, A. (2016). Test–retest: Agreement or reliability? Methodological Innovations, 9, 2059799116672875. 10.1177/2059799116672875

Bland, J. M., & Altman, D. (1986). Statistical methods for assessing agreement between two methods of clinical measurement. The Lancet, 327(8476), 307–310.

Bland, J. M., & Altman, D. G. (1999). Measuring agreement in method comparison studies. Statistical Methods in Medical Research, 8(2), 135–160.

Bogdanov, M., Renault, H., LoParco, S., Weinberg, A., & Otto, A. R. (2022). Cognitive effort exertion enhances electrophysiological responses to rewarding outcomes. Cerebral Cortex, 32(19), 4255–4270.

Braver, T. S. (2012). The variable nature of cognitive control: A dual mechanisms framework. Trends in Cognitive Sciences, 16(2), 106–113. 10.1016/j.tics.2011.12.010

Button, K. S., Ioannidis, J. P. A., Mokrysz, C., Nosek, B. A., Flint, J., Robinson, E. S. J., & Munafò, M. R. (2013). Power failure: Why small sample size undermines the reliability of neuroscience. Nature Reviews Neuroscience, 14(5), 365–376. 10.1038/nrn3475

Cavanagh, J. F., Zambrano-Vazquez, L., & Allen, J. J. (2012). Theta lingua franca: A common mid-frontal substrate for action monitoring processes. Psychophysiology, 49(2), 220–238.

Clawson, A., Clayson, P. E., & Larson, M. J. (2013). Cognitive control adjustments and conflict adaptation in major depressive disorder. Psychophysiology, 50(8), 711–721. 10.1111/psyp.12066

Clayson, P. E. (2024). The psychometric upgrade psychophysiology needs. Psychophysiology, 61(3), e14522.

Clayson, P. E., Carbine, K. A., Baldwin, S. A., & Larson, M. J. (2019). Methodological reporting behavior, sample sizes, and statistical power in studies of event-related potentials: Barriers to reproducibility and replicability. Psychophysiology, 56(11), e13437. 10.1111/psyp.13437

Cook, D. A., & Beckman, T. J. (2006). Current concepts in validity and reliability for psychometric instruments: Theory and application. The American Journal of Medicine, 119(2), 166–e7.

de Aguiar Neto, F. S., & Rosa, J. L. G. (2019). Depression biomarkers using non-invasive EEG: A review. Neuroscience & Biobehavioral Reviews, 105, 83–93. 10.1016/j.neubiorev.2019.07.021

Dell’Acqua, C., Hajcak, G., Amir, N., Santopetro, N. J., Brush, C. J., & Meyer, A. (2023). Error-related brain activity: A time-domain and time-frequency investigation in pediatric obsessive–compulsive disorder. Psychophysiology, 60(4), e14216.

Duda, J. M., Moser, A. D., Zuo, C. S., Du, F., Chen, X., Perlo, S., Richards, C. E., Nascimento, N., Ironside, M., & Crowley, D. J. (2021). Repeatability and reliability of GABA measurements with magnetic resonance spectroscopy in healthy young adults. Magnetic Resonance in Medicine, 85(5), 2359–2369.

Enkavi, A. Z., Eisenberg, I. W., Bissett, P. G., Mazza, G. L., MacKinnon, D. P., Marsch, L. A., & Poldrack, R. A. (2019). Large-scale analysis of test–retest reliabilities of self-regulation measures. Proceedings of the National Academy of Sciences, 116(12), 5472–5477. 10.1073/pnas.1818430116

Enkavi, A. Z., & Poldrack, R. A. (2021). Implications of the Lacking Relationship Between Cognitive Task and Self-report Measures for Psychiatry. Biological Psychiatry: Cognitive Neuroscience and Neuroimaging, 6(7), 670–672. 10.1016/j.bpsc.2020.06.010

Eriksen, B. A., & Eriksen, C. W. (1974). Effects of noise letters upon the identification of a target letter in a nonsearch task. Perception & Psychophysics, 16(1), 143–149. 10.3758/BF03203267

Eriksen, C. W. (1995). The flankers task and response competition: A useful tool for investigating a variety of cognitive problems. Visual Cognition, 2(2–3), 101–118.

Falkenstein, M., Hohnsbein, J., Hoormann, J., & Blanke, L. (1991). Effects of crossmodal divided attention on late ERP components. II. Error processing in choice reaction tasks. Electroencephalography and Clinical Neurophysiology, 78(6), 447–455. 10.1016/0013-4694(91)90062-9

Gehring, W. J., Goss, B., Coles, M. G. H., Meyer, D. E., & Donchin, E. (1993). A Neural System for Error Detection and Compensation. Psychological Science, 4(6), 385–390. 10.1111/j.1467-9280.1993.tb00586.x

Goschke, T. (2014). Dysfunctions of decision-making and cognitive control as transdiagnostic mechanisms of mental disorders: Advances, gaps, and needs in current research. International Journal of Methods in Psychiatric Research, 23(S1), 41–57. 10.1002/mpr.1410

Grahek, I., Everaert, J., Krebs, R. M., & Koster, E. H. (2018). Cognitive control in depression: Toward clinical models informed by cognitive neuroscience. Clinical Psychological Science, 6(4), 464–480.

Gratton, G., Coles, M. G., & Donchin, E. (1992). Optimizing the use of information: Strategic control of activation of responses. Journal of Experimental Psychology: General, 121(4), 480.

Hajcak, G., Meyer, A., & Kotov, R. (2017). Psychometrics and the neuroscience of individual differences: Internal consistency limits between-subjects effects. Journal of Abnormal Psychology, 126(6), 823.

Hedge, C., Powell, G., & Sumner, P. (2018). The reliability paradox: Why robust cognitive tasks do not produce reliable individual differences. Behavior Research Methods, 50, 1166–1186.

Heil, M., Osman, A., Wiegelmann, J., Rolke, B., & Hennighausen, E. (2000). N200 in the Eriksen-task: Inhibitory executive process? Journal of Psychophysiology, 14(4), 218.

Hopkins, W. G. (2000). Measures of reliability in sports medicine and science. Sports Medicine, 30, 1–15.

Insel, T. R. (2014). The NIMH Research Domain Criteria (RDoC) Project: Precision Medicine for Psychiatry. American Journal of Psychiatry, 171(4), 395–397. 10.1176/appi.ajp.2014.14020138

Ip, C.-T., Ganz, M., Ozenne, B., Sluth, L. B., Gram, M., Viardot, G., l’Hostis, P., Danjou, P., Knudsen, G. M., & Christensen, S. R. (2018). Pre-intervention test-retest reliability of EEG and ERP over four recording intervals. International Journal of Psychophysiology, 134, 30–43. 10.1016/j.ijpsycho.2018.09.007

Kałamała, P., Szewczyk, J., Senderecka, M., & Wodniecka, Z. (2018). Flanker task with equiprobable congruent and incongruent conditions does not elicit the conflict N2. Psychophysiology, 55(2), e12980. 10.1111/psyp.12980

Kanske, P., & Kotz, S. A. (2010). Modulation of early conflict processing: N200 responses to emotional words in a flanker task. Neuropsychologia, 48(12), 3661–3664.

Karakaş, S., Erzengin, Ö. U., & Başar, E. (2000). The genesis of human event-related responses explained through the theory of oscillatory neural assemblies. Neuroscience Letters, 285(1), 45–48.

Keil, A., Debener, S., Gratton, G., Junghöfer, M., Kappenman, E. S., Luck, S. J., Luu, P., Miller, G. A., & Yee, C. M. (2014). Committee report: Publication guidelines and recommendations for studies using electroencephalography and magnetoencephalography. Psychophysiology, 51(1), 1–21.

Kessler, R. C. (1977). The use of change scores as criteria in longitudinal survey research. Quality and Quantity, 11(1), 43–66. 10.1007/BF00143986

Koo, T. K., & Li, M. Y. (2016). A guideline of selecting and reporting intraclass correlation coefficients for reliability research. Journal of Chiropractic Medicine, 15(2), 155–163.

Kopp, B., Rist, F., & Mattler, U. (1996). N200 in the flanker task as a neurobehavioral tool for investigating executive control. Psychophysiology, 33(3), 282–294.

Laming, D. (1979). Choice reaction performance following an error. Acta Psychologica, 43(3), 199–224.

Larson, M. J., Baldwin, S. A., Good, D. A., & Fair, J. E. (2010). Temporal stability of the error-related negativity (ERN) and post-error positivity (Pe): The role of number of trials. Psychophysiology, 47(6), 1167–1171.

Lehnert, B., & Lehnert, M. B. (2015). Package ‘BlandAltmanLeh.’ *CRAN. Available Online:* https://Cran.r-Project.Org/Web/Packages/BlandAltmanLeh/BlandAltmanLeh.Pdf *(Accessed on 15 October 2016)*.

Lesh, T. A., Niendam, T. A., Minzenberg, M. J., & Carter, C. S. (2011). Cognitive control deficits in schizophrenia: Mechanisms and meaning. Neuropsychopharmacology, 36(1), 316–338.

Lin, M.-H., Davies, P. L., Stephens, J., & Gavin, W. J. (2020). Test–retest reliability of electroencephalographic measures of performance monitoring in children and adults. Developmental Neuropsychology, 45(6), 341– 366.

Luciana, M., & Collins, P. F. (2022). Neuroplasticity, the Prefrontal Cortex, and Psychopathology-Related Deviations in Cognitive Control. Annual Review of Clinical Psychology, 18 (Volume 18, 2022), 443–469. 10.1146/annurev-clinpsy-081219-111203

Luck, S. J., & Gaspelin, N. (2017). How to get statistically significant effects in any ERP experiment (and why you shouldn’t). Psychophysiology, 54(1), 146–157.

McEvoy, L., Smith, M., & Gevins, A. (2000). Test–retest reliability of cognitive EEG. Clinical Neurophysiology, 111(3), 457–463.

McLoughlin, G., Gyurkovics, M., Palmer, J., & Makeig, S. (2022). Midfrontal Theta Activity in Psychiatric Illness: An Index of Cognitive Vulnerabilities Across Disorders. Biological Psychiatry, 91(2), 173–182. 10.1016/j.biopsych.2021.08.020

McLoughlin, G., Makeig, S., & Tsuang, M. T. (2014). In search of biomarkers in psychiatry: EEG-based measures of brain function. American Journal of Medical Genetics Part B: Neuropsychiatric Genetics, 165(2), 111– 121.

McTeague, L. M., Goodkind, M. S., & Etkin, A. (2016). Transdiagnostic impairment of cognitive control in mental illness. Journal of Psychiatric Research, 83, 37–46. 10.1016/j.jpsychires.2016.08.001

McTeague, L. M., Huemer, J., Carreon, D. M., Jiang, Y., Eickhoff, S. B., & Etkin, A. (2017). Identification of Common Neural Circuit Disruptions in Cognitive Control Across Psychiatric Disorders. American Journal of Psychiatry, 174(7), 676–685. 10.1176/appi.ajp.2017.16040400

Miller, T. B., & Kane, M. (2001). The Precision of Change Scores Under Absolute and Relative Interpretations. Applied Measurement in Education, 14(4), 307–327. 10.1207/S15324818AME1404_1

Montague, P. R., Dolan, R. J., Friston, K. J., & Dayan, P. (2012). Computational psychiatry. Trends in Cognitive Sciences, 16(1), 72–80. 10.1016/j.tics.2011.11.018

Morand-Beaulieu, S., Perrault, M.-A., & Lavoie, M. E. (2021). Test-retest reliability of event-related potentials across three tasks. Journal of Psychophysiology.

Muir, A. M., Hedges-Muncy, A., Clawson, A., Carbine, K. A., & Larson, M. J. (2020). Dimensions of anxiety and depression and neurophysiological indicators of error-monitoring: Relationship with delta and theta oscillatory power and error-related negativity amplitude. Psychophysiology, 57(9), e13595. 10.1111/psyp.13595

Niv, Y. (2021). The primacy of behavioral research for understanding the brain. Behavioral Neuroscience, 135(5), 601–609. 10.1037/bne0000471

Olbrich, S., & Arns, M. (2013). EEG biomarkers in major depressive disorder: Discriminative power and prediction of treatment response. International Review of Psychiatry, 25(5), 604–618.

Olvet, D. M., & Hajcak, G. (2009). The stability of error-related brain activity with increasing trials. Psychophysiology, 46(5), 957–961. 10.1111/j.1469-8986.2009.00848.x

Paap, K. R., & Sawi, O. (2016). The role of test-retest reliability in measuring individual and group differences in executive functioning. Journal of Neuroscience Methods, 274, 81–93. 10.1016/j.jneumeth.2016.10.002

Parsons, S., Kruijt, A.-W., & Fox, E. (2019). Psychological science needs a standard practice of reporting the reliability of cognitive-behavioral measurements. Advances in Methods and Practices in Psychological Science, 2(4), 378–395.

Pathania, A., Schreiber, M., Miller, M. W., Euler, M. J., & Lohse, K. R. (2021). Exploring the reliability and sensitivity of the EEG power spectrum as a biomarker. International Journal of Psychophysiology, 160, 18–27. 10.1016/j.ijpsycho.2020.12.002

Pizzagalli, D. A., Jahn, A. L., & O’Shea, J. P. (2005). Toward an objective characterization of an anhedonic phenotype: A signal-detection approach. Biological Psychiatry, 57(4), 319–327. 10.1016/j.biopsych.2004.11.026

Pronk, T., Molenaar, D., Wiers, R. W., & Murre, J. (2022). Methods to split cognitive task data for estimating split-half reliability: A comprehensive review and systematic assessment. Psychonomic Bulletin & Review, 29(1), 44–54. 10.3758/s13423-021-01948-3

Rabbitt, P., & Rodgers, B. (1977). What does a man do after he makes an error? An analysis of response programming. The Quarterly Journal of Experimental Psychology, 29(4), 727–743.

Ramey, T., & Regier, P. S. (2019). Cognitive impairment in substance use disorders. CNS Spectrums, 24(1), 102–113. 10.1017/S1092852918001426

Riesel, A., Endrass, T., & Weinberg, A. (2021). Biomarkers of mental disorders: Psychophysiological measures as indicators of mechanisms, risk, and outcome prediction. International Journal of Psychophysiology.

Riesel, A., Weinberg, A., Endrass, T., Meyer, A., & Hajcak, G. (2013). The ERN is the ERN is the ERN? Convergent validity of error-related brain activity across different tasks. Biological Psychology, 93(3), 377–385. 10.1016/j.biopsycho.2013.04.007

Romer, A. L., & Pizzagalli, D. A. (2022). Associations Between Brain Structural Alterations, Executive Dysfunction, and General Psychopathology in a Healthy and Cross-Diagnostic Adult Patient Sample. Biological Psychiatry Global Open Science, 2(1), 17–27. 10.1016/j.bpsgos.2021.06.002

Ruchsow, M., Herrnberger, B., Wiesend, C., Grön, G., Spitzer, M., & Kiefer, M. (2004). The effect of erroneous responses on response monitoring in patients with major depressive disorder: A study with event-related potentials. Psychophysiology, 41(6), 833–840. 10.1111/j.1469-8986.2004.00237.x

Sanders, L. M., Hortobágyi, T., Balasingham, M., Van der Zee, E. A., & van Heuvelen, M. J. (2019). Psychometric properties of a flanker task in a sample of patients with dementia: A pilot study. Dementia and Geriatric Cognitive Disorders Extra, 8(3), 382–392.

Sandre, A., & Weinberg, A. (2019). Neither wrong nor right: Theta and delta power increase during performance monitoring under conditions of uncertainty. International Journal of Psychophysiology, 146, 225–239. 10.1016/j.ijpsycho.2019.09.015

Schnack, H. G., & Kahn, R. S. (2016). Detecting Neuroimaging Biomarkers for Psychiatric Disorders: Sample Size Matters. Frontiers in Psychiatry, 7. 10.3389/fpsyt.2016.00050

Shechtman, O. (2013). The coefficient of variation as an index of measurement reliability. In Methods of clinical epidemiology (pp. 39–49). Springer.

Sheehan, D. V., Lecrubier, Y., Sheehan, K. H., Amorim, P., Janavs, J., Weiller, E., Hergueta, T., Baker, R., & Dunbar, G. C. (1998). The Mini-International Neuropsychiatric Interview (MINI): The development and validation of a structured diagnostic psychiatric interview for DSM-IV and ICD-10. Journal of Clinical Psychiatry, 59(20), 22–33.

Snyder, H. R., Miyake, A., & Hankin, B. L. (2015). Advancing understanding of executive function impairments and psychopathology: Bridging the gap between clinical and cognitive approaches. Frontiers in Psychology, 6. 10.3389/fpsyg.2015.00328

Tenke, C. E., Kayser, J., Pechtel, P., Webb, C. A., Dillon, D. G., Goer, F., Murray, L., Deldin, P., Kurian, B. T., McGrath, P. J., Parsey, R., Trivedi, M., Fava, M., Weissman, M. M., McInnis, M., Abraham, K., E. Alvarenga, J., Alschuler, D. M., Cooper, C., … Bruder, G. E. (2017). Demonstrating test-retest reliability of electrophysiological measures for healthy adults in a multisite study of biomarkers of antidepressant treatment response. Psychophysiology, 54(1), 34–50. 10.1111/psyp.12758

Vanderhasselt, M.-A., Raedt, R. D., Dillon, D. G., Dutra, S. J., Brooks, N., & Pizzagalli, D. A. (2012). Decreased cognitive control in response to negative information in patients with remitted depression: An event-related potential study. Journal of Psychiatry and Neuroscience, 37(4), 250–258. 10.1503/jpn.110089

Vasey, M. W., Dalgleish, T., & Silverman, W. K. (2003). Research on information-processing factors in child and adolescent psychopathology: A critical commentary. Journal of Clinical Child and Adolescent Psychology, 32(1), 81–93.

Vaz, S., Falkmer, T., Passmore, A. E., Parsons, R., & Andreou, P. (2013). The case for using the repeatability coefficient when calculating test–retest reliability. PloS One, 8(9), e73990.

Wacker, J., Dillon, D. G., & Pizzagalli, D. A. (2009). The role of the nucleus accumbens and rostral anterior cingulate cortex in anhedonia: Integration of resting EEG, fMRI, and volumetric techniques. NeuroImage, 46(1), 327–337. 10.1016/j.neuroimage.2009.01.058

Weinberg, A., & Hajcak, G. (2011). Longer term test–retest reliability of error-related brain activity. Psychophysiology, 48(10), 1420–1425.

Weir, J. P. (2005). Quantifying test-retest reliability using the intraclass correlation coefficient and the SEM. The Journal of Strength & Conditioning Research, 19(1), 231–240.

Whitehead, P. S., Brewer, G. A., & Blais, C. (2019). Are cognitive control processes reliable? *Journal of Experimental Psychology: Learning*, Memory, and Cognition, 45(5), 765–778. 10.1037/xlm0000632

Wickham, H. (2011). Ggplot2. Wiley Interdisciplinary Reviews: Computational Statistics, 3(2), 180–185.

Williams, L., Simms, E., Clark, C., Paul, R., Rowe, D., & Gordon, E. (2005). The test-retest reliability of a standardized neurocognitive and neurophysiological test battery:“neuromarker.” International Journal of Neuroscience, 115(12), 1605–1630.

Wöstmann, N. M., Aichert, D. S., Costa, A., Rubia, K., Möller, H.-J., & Ettinger, U. (2013). Reliability and plasticity of response inhibition and interference control. Brain and Cognition, 81(1), 82–94. 10.1016/j.bandc.2012.09.010

Xu, I., Passell, E., Strong, R. W., Grinspoon, E., Jung, L., Wilmer, J. B., & Germine, L. T. (2024). No Evidence of Reliability Across 36 Variations of the Emotional Dot-Probe Task in 9,600 Participants. Clinical Psychological Science, 21677026241253826.

